# Multimodal MRI Clustering Reveals an Elevated Free-Water Biotype in Autism with Increased Social Symptoms

**DOI:** 10.64898/2026.09.25.754196

**Authors:** Faye McKenna, Kimberly J. Hemmerling, Shounak Nandi, Sophie Molholm, David C. Zhu

## Abstract

**Background:** Autism spectrum disorder (ASD) is neurobiologically heterogeneous, and group comparisons may mask effects confined to subgroups. We tested whether distinct biotypes exist within ASD using unsupervised clustering of multimodal MRI.

**Methods:** We analyzed 149 participants with ASD and 129 typically developing (TD) controls from the Autism Brain Imaging Data Exchange II (6 sites; age 5 to 64 years). Site-harmonized features comprised free water (FW), tissue anisotropy and diffusivity, diffusion tensor image analysis along the perivascular space (DTI-ALPS), cerebrospinal fluid (CSF) volumes and exploratory resting-state regional homogeneity. ASD participants were clustered with k-means using global (7 features) and regional (17 features) sets, with robustness tested by permutation and two further algorithms. TD participants were projected post hoc to test specificity. A composite imaging score was related to Social Responsiveness Scale (SRS) subscales, symptom severity and IQ using partial correlations with false discovery rate correction.

**Results:** Clustering identified an elevated free-water (EFW) biotype comprising approximately 10% of ASD participants, defined by markedly increased FW (d = +4.91 versus remaining ASD participants), reduced tissue anisotropy, reduced DTI-ALPS and enlarged ventricles. It was recovered across feature sets and algorithms (permutation p < .0001) and departed from TD norms, unlike the remaining ASD participants. The composite score correlated with social-perceptual and communicative SRS subscales, most strongly social awareness (r = +0.253, q = .026), but not with symptom severity or IQ. FW related to social motivation in children and adolescents (r = +0.287, q = .037) but not adults.

**Limitations:** Each person was scanned once, so we cannot tell whether these brain differences cause social difficulties or how they change with age. The subgroup was small, clinical data were incomplete, the sample was mostly male with a wide age range, and the findings await confirmation in an independent group. Free water can reflect inflammation, swelling or tissue loss, so its biological meaning remains uncertain.

**Conclusions:** We identify an ASD subgroup defined by extracellular fluid excess and altered white matter that is associated with greater social difficulties. If replicated, this signature may help stratify individuals with ASD in mechanistic studies and clinical trials.

## Background

Autism spectrum disorder (ASD) is among the most neurobiologically heterogeneous of neurodevelopmental conditions, with substantial variability in etiology, brain structure and symptom expression [1,2]. This heterogeneity has impeded the identification of reproducible imaging biomarkers and the development of targeted interventions [3]. One strategy for parsing this heterogeneity is data-driven biotyping, in which unsupervised clustering is applied to neuroimaging and/or peripheral biomarker features to identify biologically distinct subgroups [4]. This approach has demonstrated clinical relevance in schizophrenia and depression [5–7], and has identified subtypes with distinct progression and treatment response in multiple sclerosis and neurodegenerative disease [8–10]. It has also been applied in ASD using structural morphometry or functional connectivity [2,11–14]. Diffusion-based biotyping is more recent, with isotropic diffusion metrics used to identify Parkinson’s disease subtypes [15,16]. To our knowledge, no study in any condition has used unsupervised clustering on a combined panel indexing extracellular fluid, periventricular diffusivity, white matter microstructure and local functional coherence to derive biotypes.

The first of these, extracellular fluid, is indexed by the free water (FW) fraction. Derived from diffusion MRI modeling, FW separates extracellular from tissue-bound water and provides a sensitive marker of neuroinflammation, axonal loss and extracellular fluid accumulation [17–19]. The same free water imaging (FWI) model returns tissue mean diffusivity (MDt) and tissue fractional anisotropy (FAt), which are conventional MD and FA with the free water contribution removed. Because the free water and tissue compartments are estimated jointly, the pattern across measures is informative: elevated FW accompanied by elevated MDt and reduced FAt indicates disruption within the tissue compartment itself rather than displacement of tissue by fluid, a distinction conventional diffusion tensor imaging (DTI) cannot make [20]. Animal work ties FW to specific cellular events: in rats, induced neuroinflammation and demyelination raise diffusivity in proportion to microglial and astrocytic activation, validated against histology [21]; in mice, interferon-gamma overexpression raises FW via a defined signaling cascade [22]; and maternal immune activation, the principal rodent model of ASD risk, elevates FW in adult offspring [23]. Consistent with this, thus far FW elevation has been documented across the lifespan in ASD, in cortico-basal ganglia tracts in children, frontal white matter in adults, and the hippocampus in older adults [24–27].

Fluid excess in ASD is not confined to brain tissue; it also accumulates around the brain in the CSF spaces, where some of the earliest detectable evidence appears. Increased extra-axial CSF is detectable at 6 months in infants later diagnosed with ASD, precedes behavioral symptoms, and predicts diagnosis and severity at 36 months [28]; however, it may not persist, as one study from middle childhood through adulthood found CSF excess no longer elevated [29]. On this basis, CSF excess has been proposed to mark a distinct, developmentally related autism subtype [30]. Together these point to excess fluid in ASD both within and around the brain. That it appears prominent early but not later suggests it is carried by a subset of individuals and would be diluted by averaging over the whole diagnosis or a wide age range, motivating approaches that resolve which individuals carry the signal rather than testing ASD as a single population.

Extracellular fluid excess in ASD raises the question of whether brain fluid regulation is altered, and, if so, where its structural signature would be most detectable. Periventricular white matter is a plausible candidate region since perivascular spaces in this territory support fluid exchange between CSF, perivascular, and interstitial compartments [31,32], and the region is among the most vulnerable to fluid accumulation and white matter injury [33]. Since altered regulation would reshape the extracellular geometry and perivascular conditions of this tissue, a measure sensitive to those properties can serve as its readout. The diffusion tensor image analysis along the perivascular space (DTI-ALPS) index samples diffusivity along versus perpendicular to the dominant fiber orientation adjacent to the lateral ventricular body, and because its value reflects white matter architecture, fiber orientation, extracellular geometry, and perivascular conditions jointly, it is an indirect but promising structural biomarker of the periventricular tissue environment [34,35]. Notably, reduced ALPS has been reported in children with ASD [36–38] and in aging and psychiatric disorders more broadly [39,40]. These findings support the inclusion of DTI-ALPS alongside free water imaging and CSF metrics within a multimodal clustering framework.

Extracellular fluid, tissue microstructure, CSF volume and periventricular diffusion reflect related aspects of a shared fluid regulation and clearance biology, but they have been studied largely in isolation, and each has yielded an inconsistent literature in ASD. Such inconsistency is the expected signature of an effect driven by subsets of individuals rather than by the diagnostic group as a whole, and it is not resolvable by case-control comparison, which masks subgroup heterogeneity. In the current study, we applied unsupervised clustering to multimodal MRI features acquired in 278 participants from Autism Brain Imaging Data Exchange II (ABIDE II) [41] (149 with ASD and 129 TD controls; 6 sites; age 5 to 64 years), harmonized across sites using neuroCombat [42,43] (Additional file 1: Tables S1, S2). We also included one exploratory functional feature, regional homogeneity (ReHo), the temporal coherence of the resting-state BOLD signal within a local voxel neighborhood [44]. The feature space thus spans microstructural diffusion, macroscale CSF volumes, and hemodynamic coherence; combining modalities this way can reveal variation only partially captured by any one [45,46], consistent with biotyping that combines measures across biological levels [47].

Two feature sets were evaluated in sequence: a primary global set (n = 7) of whole-brain averages of these metrics, and a secondary regional set (n = 17) of anterior, medial and posterior positional means. This subdivision is anatomically motivated: periventricular clearance is regionally organized rather than uniform, with the deep medullary veins and their perivascular spaces converging on distinct subependymal territories along the anteroposterior axis [48], and periventricular vulnerability to fluid accumulation and white matter injury varies along the same axis [49]. Because DTI-ALPS is defined at a fixed periventricular location adjacent to the lateral ventricle body, where the medullary-vein perivascular spaces run orthogonal to the projection and association fibers [35], the regional parcellation was built to match its ventricular-body boundaries, so that every marker except the ventricles is sampled within the same anatomical zones where the ALPS geometry remains valid (Figure 1A-B). This lets us ask whether the biotype depends on the regional organization of periventricular drainage and vulnerability, or is recoverable from whole-brain averages alone.

**Figure 1.**
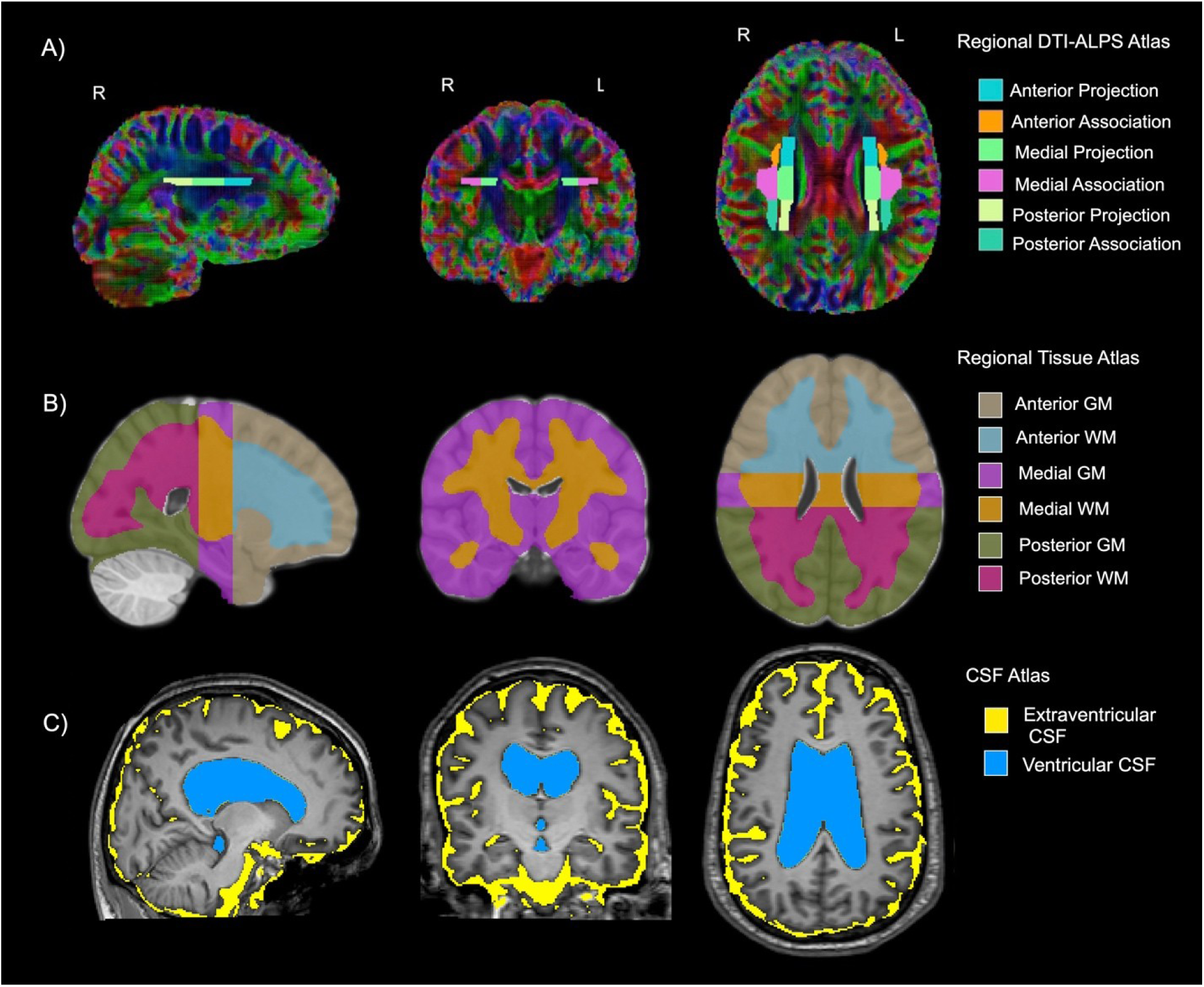
Regional neuroimaging atlases used for DTI-ALPS, white matter/gray matter segmentation, and CSF analyses. **(A) Regional DTI-ALPS Atlas.** Bilateral projection and association fiber ROIs displayed on a color-coded FA map in sagittal (left), coronal (center), and axial (right) views, divided into three rostrocaudal segments corresponding to the anterior, medial, and posterior ventricular body. Projection ROIs (corona radiata) and association ROIs (superior longitudinal fasciculus) are shown for each segment by color coding: cyan = anterior projection; **orange** = anterior association; **green** = medial projection; **pink** = medial association; **yellow-green** = posterior projection; **teal** = posterior association. See Methods for full ROI derivation and ventricle-fitting procedure. **(B) Rostrocaudal Regional Atlas for FWI and ReHo Analysis.** Custom whole-brain atlas dividing the brain into anterior, medial, and posterior thirds along the rostrocaudal axis — defined to match the DTI-ALPS regional boundaries — each subdivided into gray and white matter compartments, shown in sagittal (left), coronal (center), and axial (right) views. Brain regions are color coded. Colors denote: **tan** = anterior gray matter; **blue** = anterior white matter; **purple** = medial gray matter; **orange** = medial white matter; **green** = posterior gray matter; **pink** = posterior white matter. **(C) CSF** volume estimation in the **yellow** = extraventricular and **blue** = ventricular space.

We predicted that this feature space would contain neurobiologically distinct subgroups within ASD rather than continuous variation across the diagnostic group. We further tested whether any such subgroup is specific to ASD, by projecting TD participants post-hoc into the ASD-derived cluster space. Finally, we derived a composite imaging score from the same features and tested whether it related to social functioning, cognitive ability or symptom severity, and whether associations related to developmental stage across the wide age range.

## Methods

See Additional file 1 for more information.

### Participants

Data were drawn from the ABIDE II dataset, an open-access multisite repository [50]. Following quality control, 278 participants who had diffusion, anatomical and functional MRI data from six sites were selected for analyses: 149 ASD (mean age = 16.5 ± 13.5 years; 128 M / 18 F; IQ = 103.5 ± 17.6) and 129 TD (mean age = 21.3 ± 14.5 years; 104 M / 24 F; IQ = 112.2 ± 14.4).

Sites: BNI, NYU1, NYU2, IP, SDSU and TCD [50]. ASD diagnosis was confirmed per site protocol using Autism Diagnostic Observation Schedule–Generic (ADOS-G) [51] and/or Autism Diagnostic Interview-Revised (ADI-R) [52]. See Table 1 for demographic and clinical characteristics.

**Table 1.** Participant Demographics and Clinical Characteristics Values are mean ± SD unless otherwise noted. d = Cohen’s d (ASD − TD). ASD and TD groups were not age-matched by design in ABIDE II; the age difference is therefore reported as a sample characteristic, while age and sex were covaried in all subsequent imaging analyses. SRS = Social Responsiveness Scale (parent- or self-report, T-scores); ADOS = Autism Diagnostic Observation Schedule; BDI-II = Beck Depression Inventory-II. ^a^Age and sex: ASD n = 146, TD n = 128; four participants lacked demographic data and were excluded from analyses using age or sex as covariates. ^b^IQ: ASD n = 142 (FIQ), 113 (VIQ), 116 (PIQ); TD n = 102 (FIQ), 68 (VIQ), 94 (PIQ). ^c^BDI-II: ASD n = 43, TD n = 41. ^d^SRS: ASD n = 125, TD n = 94. ^e^ADOS-G: ASD n = 88; not administered to TD group. **p < .01.

| *Variable* | *ASD (n = 149)* | *TD (n = 129)* | *Statistic* | *p* | *d* |
| --- | --- | --- | --- | --- | --- |
| *Age (years)<sup>a</sup>* | 16.5 ± 13.5 | 21.3 ± 14.5 | t = -2.86 | .005** | -0.35 |
| *Sex (M/F)<sup>a</sup>* | 128 / 18 | 104 / 24 | χ<sup>2</sup> = 1.70 | .192 | — |
| *Full-Scale IQ<sup>b</sup>* | 103.5 ± 17.6 | 112.2 ± 14.4<sup>s</sup> | t = -4.10 | <.001** | -0.54 |
| *Verbal IQ<sup>b</sup>* | 103.0 ± 18.0 | 113.5 ± 14.9 | t = -4.07 | <.001** | -0.62 |
| *Performance IQ<sup>b</sup>* | 102.7 ± 19.8 | 110.5 ± 14.2 | t = -3.24 | .001** | -0.45 |
| *BDI-II Total<sup>c</sup>* | 9.8 ± 8.9 | 4.8 ± 6.0 | t = 2.99 | .004** | +0.65 |
| *SRS Total T<sup>d</sup>* | 77.7 ± 12.5 | 45.7 ± 9.6 | t = 20.57 | <.001** | +2.86 |
| *SRS Awareness T<sup>d</sup>* | 68.6 ± 12.7 | 44.9 ± 9.4 | t = 15.30 | <.001** | +2.13 |
| *SRS Cognition T<sup>d</sup>* | 74.7 ± 13.2 | 43.8 ± 6.8 | t = 20.65 | <.001** | +2.93 |
| *SRS Communication T<sup>d</sup>* | 74.9 ± 13.3 | 44.0 ± 6.2 | t = 21.54 | <.001** | +3.07 |
| *SRS Motivation T<sup>d</sup>* | 72.0 ± 12.9 | 47.2 ± 7.5 | t = 16.61 | <.001** | +2.35 |
| *SRS Mannerisms T<sup>d</sup>* | 79.1 ± 16.1 | 45.9 ± 6.0 | t = 18.99 | <.001** | +2.73 |
| *ADOS Total<sup>e</sup>* | 11.0 ± 4.0 | — | — | — | — |
| ADOS<br>Communication<sup>e</sup> | 3.6 ± 1.8 | — | — | — | — |
| ADOS Social<sup>e</sup> | 7.4 ± 2.6 | — | — | — | — |

Multi-site harmonization was performed using neuroCombat [42,43] (empirical Bayes; parametric; mean and variance correction), with ASD/TD diagnosis preserved as a biological covariate.

### Cognitive and Clinical Assessment

ASD symptom severity was assessed using the ADOS-G [51], which includes standardized observer-rated domains of social affect and communication. Dimensional social responsiveness was assessed via the Social Responsiveness Scale (SRS) [53] T-scores across five subscales: social awareness, cognition, communication, motivation, and restricted/repetitive behavior (mannerisms). General cognitive ability was assessed via full-scale IQ (ASD n = 142; TD n = 102) using site-specific batteries [54–56].

### MRI Acquisition

Diffusion MRI data were acquired across six sites on 3T scanners with site-specific protocols (see ABIDE II documentation) [50]. T1-weighted and resting-state fMRI data were acquired at all sites. All data passed ABIDE II quality control, with additional checks prior to harmonization (see Additional file 1).

### Regional Parcellation

All T1-weighted images were processed through the FSL’s segmentation pipeline [57]. The white and gray matter surrounding the lateral ventricles was parcellated into anterior, medial, and posterior zones whose boundaries were taken directly from the DTI-ALPS ventricular-body ROIs (Figure 1A). The parcellation was therefore anchored to the anatomy at which DTI-ALPS is defined, so that every marker is sampled within the same periventricular zones and applied to FWI and ReHo metrics (Figure 1B). Ventricular and extraventricular CSF compartments were derived from FSL FAST partial volume estimates (Figure 1C) [58].

### Diffusion Preprocessing

Preprocessing used MRtrix3 [59], FSL [60] and Synb0-DisCo [61]: denoising, Gibbs-ringing, susceptibility, eddy-current and motion correction (with outlier replacement), and bias-field correction. Tensors were fit with FSL dtifit after brain extraction [60]. All diffusion data was registered to MNI152 [62] space for regional parcellation.

#### DTI-ALPS

The DTI-ALPS index was computed from tensor component maps (Dxx, Dyy, Dzz) using projection and association fiber ROIs from the JHU atlas at the lateral ventricular body level following previous work [35,63]. ROIs were restricted to white matter voxels with fractional anisotropy (FA) > 0.2 to exclude partial volume effects at tissue boundaries following previous work [63]. ROIs were subsequently segmented into anterior, medial, and posterior thirds, yielding regional sub-indices (aALPS, mALPS, pALPS) (Figure 1A). Bilateral regional indices were averaged, and global DTI-ALPS (gALPS) was computed as the mean across all regions.

### Free Water Imaging

Free water fraction and free-water-corrected tensor metrics were estimated using the bi-tensor free water elimination (FWE) model of Pasternak et al. [20] (Additional file 1: Figure S1A); the latter comprised free-water-corrected fractional anisotropy (FAt), mean diffusivity (MDt), axial diffusivity (ADt), and radial diffusivity (RDt), with the subscript t denoting the tissue compartment. Estimation used an MD-based initialization with the tissue MD prior calibrated to this cohort’s deep white matter and Beltrami spatial regularization (Additional file 1: Figure S1B) [64].

### Resting-State fMRI: Regional Homogeneity

Resting-state fMRI preprocessing was performed using FSL (v6.0.7) [60] and AFNI (v26.2.00) [65]. Steps included: slice-timing correction, head motion correction, functional-to-structural registration, structural-to-MNI registration, tissue segmentation, nuisance regression of white matter signal, CSF signal, and 24 motion parameters [66], and temporal bandpass filtering (0.01-0.10 Hz). Regional homogeneity (ReHo) was computed with a 27-voxel neighborhood using AFNI 3dReHo [44] (ALFF and fALFF also computed, later excluded). Maps were z-scored within each brain mask and warped to MNI space.

### Neuroimaging Feature Sets

Candidate neuroimaging features included global or regional DTI-ALPS, FAt, MDt, RDt, ADt, FW, ALFF, fALFF, ReHo, and CSF volumes. Feature selection was guided by collinearity analysis and systematic drop-one and greedy forward selection to optimize cluster separation (Additional file 1: Tables S3-S6). The global set (n = 7) comprised: gDTI-ALPS, FW global mean, MDt global mean, FAt WM total, ventricle vol/TIV, extraventricular CSF/TIV, and ReHo GM mean. The secondary regional set (n = 17) incorporated anterior/medial/posterior positional means across ALPS, FW, MDt, FAt WM, and ReHo GM, plus the two CSF/TIV measures. CSF volumes were TIV-normalized and winsorized at ±3 SD (Additional file 1). ALFF and fALFF were evaluated but excluded at feature selection (Additional file 1: Table S2).

### Dimensionality Reduction

Each feature matrix was z-scored using ASD-derived means and standard deviations, and principal component analysis was fit on ASD participants only. TD participants were then projected into this ASD-defined component space using the same scaling and PCA parameters. Components explaining ≥ 80% of variance were retained: 5 PCs (global, 87.4% variance) and 8 PCs (regional, 83.2% variance). Age and sex were not residualized beforehand, to preserve biologically relevant variance across the wide age range. Because this raises the possibility that the resulting subgroups reflect demographic rather than neurobiological structure, we repeated the clustering to check with age and sex included directly as additional features, down-weighted to 0.5 relative to the z-scored imaging features so that they could influence the solution without dominating it. Demographic confounds were additionally controlled at inference via ANCOVA.

### Neuroimaging Clustering

The primary analysis used k-means (50 initializations × 200 random seeds). Cluster number was selected by average silhouette across k = 2 to 6 (Additional file 1: Figure S2); k = 2 was optimal. Non-random structure was confirmed by permutation testing (n = 2,000). To verify that this solution was not an artifact of the algorithm, two additional methods were applied to the same features, Gaussian mixture models (GMM; 20 initializations, k selected by BIC) and hierarchical Ward clustering (cut at the optimal silhouette). Agreement across algorithms and feature sets was quantified by the adjusted Rand index (ARI). Because biotypes were defined by clustering on these features, subsequent between-biotype comparisons are descriptive rather than independent hypothesis tests and are reported as such.

### Dimensional Imaging-Clinical Analyses

To complement the binary biotype classification, a composite imaging score (PC1) was derived from the same 17 regional features used in the primary clustering analysis. PCA was fit on z-scored features (missing values imputed to feature-wise ASD medians) across all 149 ASD participants; PC1 explained 28.9% of total variance.

PC1 was related to SRS subscale T-scores, IQ standard scores and ADOS-G total using partial correlations, in the subset of ASD participants with available clinical data (SRS n = 125, ADOS-G n = 88, IQ n = 142). Age and sex were removed by OLS residualization: PC1 and each clinical subscale were separately regressed on age and sex, and Pearson r was computed between the residuals. The Benjamini-Hochberg (BH) false discovery rate (FDR) [67] procedure was applied within each subscale family.

To test whether PC1 and global mean free water (the feature most strongly differentiating the biotypes) capture distinct clinical variance, both were entered as simultaneous predictors in multiple regression models, run separately for each clinical scale family with age and sex as covariates and all predictors z-scored. The BH procedure [67] was applied across outcomes within each scale family.

### Age-Stratified FW-Cognition Associations

Given the wide age range (5-64 years), analyses were repeated a priori within children/adolescents (< 18 years) and adults (≥ 18 years). FW-SRS, ADOS, and IQ partial correlations were recomputed within each stratum (age retained as covariate), and an ANCOVA compared cluster groups on each subscale (covarying age and sex), with partial Cohen’s d = t × sqrt[(n1 + n2)/(n1 × n2)].

## Results

### Demographic and Group Differences

ASD and TD participants did not differ on sex (χZ = 1.70, p = .192). Groups differed modestly on age (d = −0.35, p = .005) and FIQ (d = −0.54, p < .001), reflecting ABIDE II’s broad age range and mixed-site composition. ASD showed markedly elevated SRS scores (all d > +2.13, all p < .001) and elevated BDI-II (d = +0.65, p = .004; Table 1). Within the ASD sample, the two clusters did not differ significantly on age, sex, or FIQ in either analysis (all p > .08). Full ASD-versus-TD comparisons for all neuroimaging markers are reported in Additional file 1: Table S8.

### Cluster Validity

The global feature set served as the primary analysis and the regional set as a secondary test of regional specificity. Both converged on k = 2 as the optimal solution by average silhouette across all three algorithms (Additional file 1: Table S6; Figure 2). Global k-means: silhouette = 0.396, permutation p < .0001 (null: −0.002 ± 0.060). Regional k-means: silhouette = 0.391, permutation p < .0001 (null: −0.003 ± 0.058). GMM selected k = 2 (global, ARI = 0.954) and k = 3 (regional, ARI = 0.615); hierarchical Ward gave k = 2 in both analyses (ARI = 0.906 global, 1.000 regional). Convergence across algorithms and significant permutation p-values confirm the two-cluster solution is robust.

**Figure 2.**
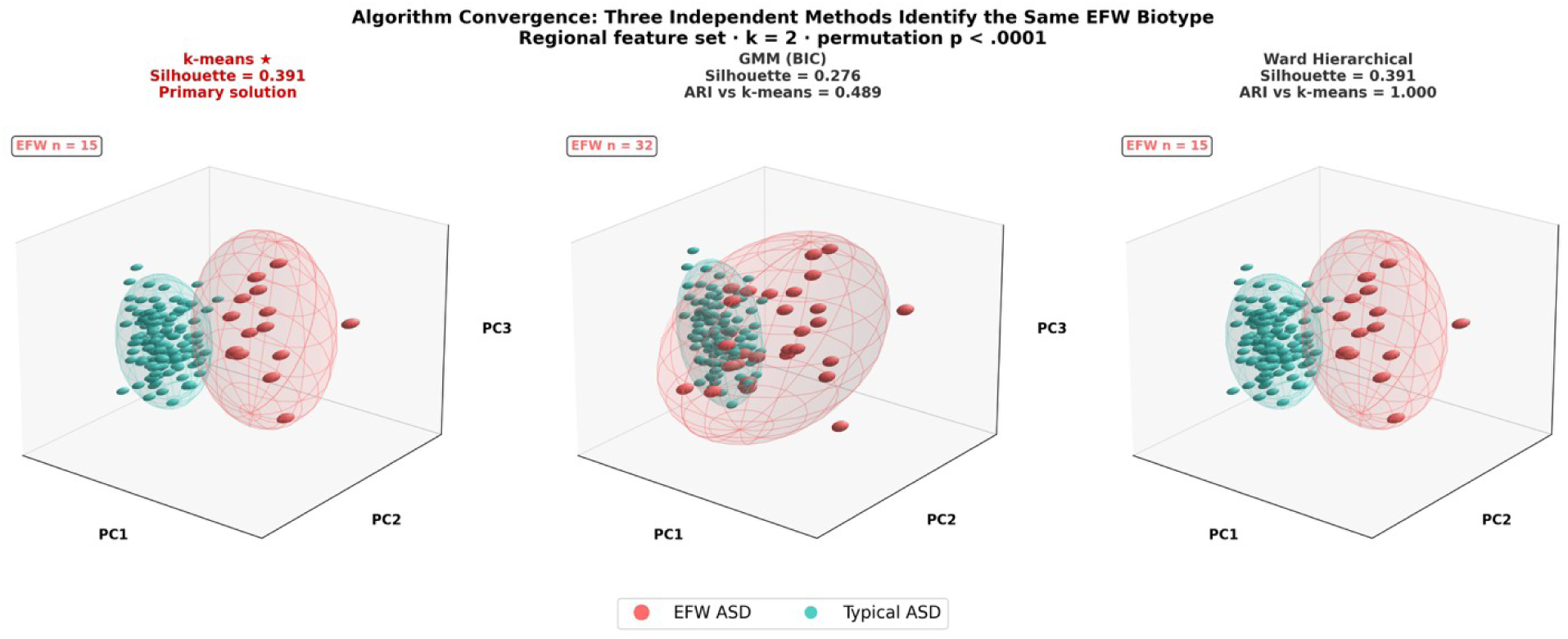
Algorithm convergence across three independent clustering methods (regional feature set, k = 2). Each panel shows all ASD subjects in the same PCA visualization space (PC1 × PC2 × PC3), colored by cluster assignment from **(A)** k-means (primary solution; 50 initializations × 200 random seeds; silhouette = 0.391; permutation p < .0001), **(B)** Gaussian Mixture Model (20 initializations; k selected by BIC; silhouette = 0.339; ARI vs k-means = 0.615), and **(C)** Ward hierarchical linkage (cut at k = 2; silhouette = 0.391; ARI vs k-means = 1.000). Salmon spheres = EFW ASD; teal spheres = non-EFW ASD; colored wireframe ellipsoids = 90% confidence regions. k-means and Ward solutions are identical (ARI = 1.000), confirming the EFW biotype is robust to algorithm choice. ARI = Adjusted Rand Index.

Including age and sex as down-weighted clustering features left assignments essentially unchanged (global ARI = 0.954; regional ARI = 1.000); the two clusters did not differ in age, and the minority cluster was not concentrated in older participants (Additional file 1: Section S3.4)

### The Elevated Free-Water (EFW) Biotype

One cluster was distinguished primarily by elevated free water and is hereafter termed the Elevated Free-Water (EFW) biotype (n = 14; 9.4% of ASD). Relative to the non-EFW biotype, it showed markedly elevated global mean FW and MDt, reduced FAt, and reduced DTI-ALPS (d = −0.60*). Extraventricular CSF/TIV and ReHo GM did not differ between biotypes (both p > .26; Table 2, Figure 3). Because the biotypes were defined by clustering on these features, these within-cluster effect sizes are descriptive of the partition rather than independent tests. To assess departure from normative values instead, both biotypes were compared to the TD group in ANCOVA (age and sex covaried). The non-EFW biotype was indistinguishable from TD in global mean FW (0.343 ± 0.018 vs 0.346 ± 0.024), whereas the EFW biotype markedly exceeded it (0.447 ± 0.039), placing it well outside the typically developing range. Notably, the whole-sample ASD-versus-TD difference in free water was modest (d = +0.31) and comparable to the group-level white-matter increase reported in schizophrenia (meta-analytic Hedges’ g = 0.38) [68], yet in this cohort it was carried almost entirely by the EFW subgroup, consistent with a subgroup effect diluted in whole-group case-control comparisons.

**Table 2.** Neuroimaging Feature Differences: EFW Biotype vs Non-EFW ASD Values are mean ± SD (ComBat site-harmonized); partial Cohen’s d from ANCOVA with age and sex covaried. CSF volume features additionally covaried for TIV. The DTI-ALPS medial effect was confirmed with free-water-corrected ALPS, indicating the reduction is not attributable to elevated extracellular water alone. ᵃCovaried for TIV. **p < .01; *p < .05; †p < .10.

| *Feature* | *EFW mean ± SD* | *Non-EFW mean ± SD* | *d (partial)* | *p* |
| --- | --- | --- | --- | --- |
| ***Global analysis — EFW biotype (n = 14 EFW, n = 135 non-EFW ASD)*** |  |  |  |  |
| *FW global mean* | 0.447 ± 0.039 | 0.343 ± 0.018 | +4.91** | <.001 |
| *MDt global mean (mm<sup>2</sup>/s)* | 0.00081 ± 0.00010 | 0.00063 ± 0.00003 | +2.87** | <.001 |
| *Ventricle vol/TIV<sup>a</sup>* | 0.0637 ± 0.0090 | 0.0556 ± 0.0088 | +1.03** | .001 |
| *FAt WM total* | 0.337 ± 0.045 | 0.364 ± 0.032 | −0.96** | .001 |
| *DTI-ALPS global* | 1.075 ± 0.104 | 1.148 ± 0.093 | −0.60* | .040 |
| *Extravent. CSF/TIV<sup>a</sup>* | 0.0432 ± 0.0055 | 0.0446 ± 0.0060 | −0.33 | .260 |
| *ReHo GM mean* | 0.110 ± 0.108 | 0.131 ± 0.084 | −0.23 | .426 |
| ***Regional analysis — EFW biotype (n = 15 EFW, n = 134 non-EFW ASD)*** |  |  |  |  |
| *FW posterior mean* | 0.485 ± 0.068 | 0.358 ± 0.028 | +3.56** | <.001 |
| *FW medial mean* | 0.445 ± 0.070 | 0.344 ± 0.018 | +3.48** | <.001 |
| *FW anterior mean* | 0.404 ± 0.072 | 0.327 ± 0.023 | +2.47** | <.001 |
| *MDt medial mean* | 0.00074 ± 0.00011 | 0.00062 ± 0.00004 | +2.47** | <.001 |
| *MDt posterior mean* | 0.00085 ± 0.00016 | 0.00070 ± 0.00007 | +2.21** | <.001 |
| *MDt anterior mean* | 0.00070 ± 0.00010 | 0.00061 ± 0.00004 | +1.59** | <.001 |
| *FAt WM posterior* | 0.330 ± 0.048 | 0.360 ± 0.033 | −1.05** | .001 |
| *FAt WM anterior* | 0.331 ± 0.051 | 0.354 ± 0.037 | −0.76** | .008 |
| *FAt WM medial* | 0.357 ± 0.044 | 0.374 ± 0.031 | −0.71* | .013 |
| *ALPS medial* | 1.086 ± 0.142 | 1.193 ± 0.121 | −0.79** | .006 |
| *ALPS posterior* | 1.043 ± 0.128 | 1.101 ± 0.116 | −0.41 | .150 |
| *ALPS anterior* | 1.100 ± 0.105 | 1.147 ± 0.111 | −0.37 | .195 |
| *Ventricle vol/TIV<sup>a</sup>* | 0.0668 ± 0.0093 | 0.0556 ± 0.0086 | +0.94** | .001 |
| *Extravent. CSF/TIV<sup>a</sup>* | 0.0437 ± 0.0055 | 0.0446 ± 0.0060 | -0.19 | .505 |
| *ReHo GM posterior* | 0.190 ± 0.255 | 0.310 ± 0.199 | -0.55<sup>†</sup> | .053 |
| *ReHo GM medial* | -0.048 ± 0.055 | -0.077 ± 0.065 | +0.44 | .124 |
| *ReHo GM anterior* | 0.224 ± 0.186 | 0.234 ± 0.176 | -0.09 | .746 |

Values are mean ± SD (ComBat site-harmonized); partial Cohen's d from ANCOVA with age and sex covaried. CSF volume features additionally covaried for TIV. The DTI-ALPS medial effect was confirmed with free-water-corrected ALPS, indicating the reduction is not attributable to elevated extracellular water alone. <sup>a</sup>Covaried for TIV. \*\*p < .01; \*p < .05; <sup>†</sup>p < .10.

**Figure 3.**
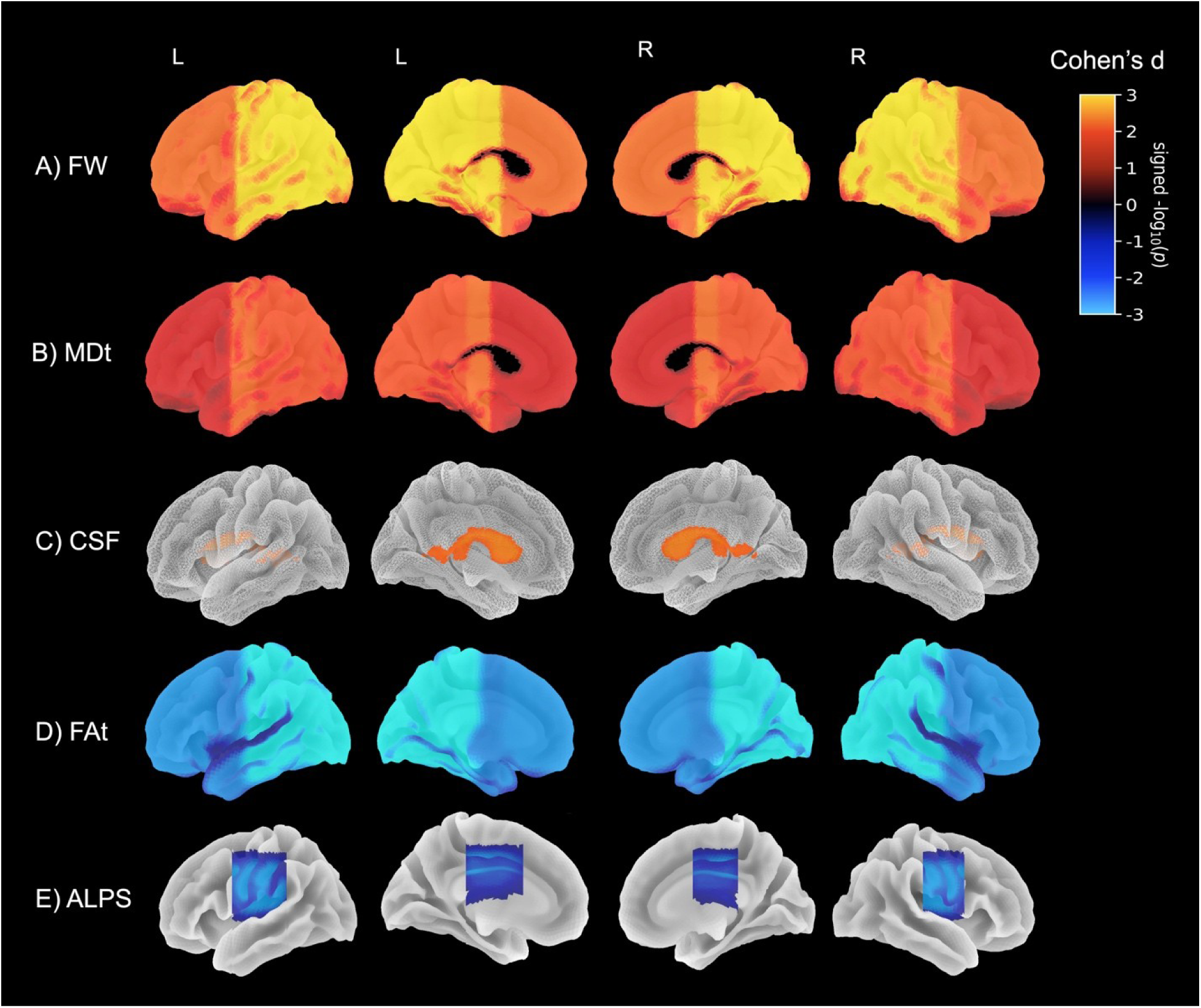
Neuroimaging profile of the elevated free-water biotype in ASD. Surface and slice renderings of neuroimaging effect sizes (partial Cohen’s d, ANCOVA age and sex covaried) for the elevated free-water (EFW) biotype versus non-EFW ASD across five imaging markers. (A) Free water fraction (FW) is markedly elevated across all anterior, medial, and posterior regions (d = +2.47 to +4.91). (B) Tissue mean diffusivity (MDt) shows uniform elevation across all A/M/P positions (d = +1.59 to +2.87). (C) Ventricular volume normalized to total intracranial volume (Vent/TIV) is enlarged bilaterally (d = +0.94 to +1.03), consistent with ex-vacuo dilatation or primary CSF production abnormalities. (D) Free-water-corrected fractional anisotropy of white matter (FAt WM) is reduced across all positions, with posterior showing the largest effect (d = −1.05**). (E) DTI-ALPS index shows a medial-predominant reduction (d = −0.79**, p = .006) at the lateral ventricular body level, while anterior and posterior positions are non-significant. Warm colors (red-yellow) indicate elevation in the EFW biotype; cool colors (blue-cyan) indicate reduction. Color intensity reflects −log₁₀(p). EFW = elevated free-water biotype; FW = free water fraction; MDt = tissue mean diffusivity; FAt WM = free-water-corrected white matter fractional anisotropy; DTI-ALPS = diffusion tensor image analysis along the perivascular space; A/M/P = anterior/ medial /posterior.

#### Regional analysis

The same subgroup emerged in the regional feature set as the EFW biotype (n = 15; 10.1% of ASD). It showed elevated FW at all three positions, with a posterior-to-anterior gradient (posterior d = +3.56**, medial d = +3.48**, anterior d = +2.47**), elevated MDt, most pronounced medially (medial d = +2.47**, posterior d = +2.21**, anterior d = +1.59**), and consistent FAt WM reductions (posterior d = −1.05**, anterior d = −0.76**, medial d = −0.71*). As in the global solution, these within-biotype effect sizes describe the clustering partition rather than constituting independent tests. DTI-ALPS showed a medial-predominant reduction: ALPS medial was significantly reduced (d = −0.79**, p = .006), whereas ALPS posterior (d = −0.41, p = .150) and ALPS anterior (d = −0.37, p = .195) did not, consistent with the ALPS decrease in the global solution (d = −0.60). Ventricular volume/TIV was enlarged (d = +0.94**, TIV-covaried), while ReHo GM posterior showed only a marginal reduction (d = −0.55, p = .053; Figure 3).

Across both feature sets, the same EFW subgroup emerged, with concordant elevations in FW and MDt and reductions in FAt and ALPS, indicating the biotype is robust to whether whole-brain or regionally resolved features are used. Two observations indicate the ALPS reduction is not merely a byproduct of diffuse extracellular fluid excess: it was confined to the medial region whereas FW was elevated at all positions, and the medial reduction persisted after free-water correction (fwi-ALPS medial: d = −0.62, p = .030), indicating it is not attributable to extracellular water contaminating the tensor.

### Cross-Analysis Convergence

The ARI between global and regional biotype assignments was 0.954. All 14 global EFW subjects were also classified as regional EFW; the additional regional-only subject represents an intermediate case captured by spatial information in the regional feature set but not by global averages alone. That the same subgroup is recovered both from whole-brain averages and from regional features indicates the biotype does not depend on the regional parcellation; the secondary regional analysis instead identifies which markers carry regional specificity, most notably DTI-ALPS.

### TD Enrichment and Neurobiological Specificity

The ASD proportion within the EFW cluster (68-70%) exceeds the overall sample ASD proportion (54%). Because the enrichment tests were themselves non-significant, this pattern is suggestive of, but does not establish, neurobiological specificity of the FW elevation; the presence of several TD participants within the cluster is equally consistent with a fluid phenotype that is expressed, less commonly, outside ASD; these projected TD participants showed an intermediate imaging profile (Additional file 1: Table S7).

### Dimensional Imaging-Clinical Associations

PC1 explained 28.9% of variance in the 17-feature regional set (loadings in Figure 4B, Additional file 1: Table S10). It was dominated by positive MDt loadings (medial +0.384, anterior +0.366) and negative FAt loadings across all three positions (−0.327 to −0.361), with moderate positive FW contributions (+0.22 to +0.29) and a moderate negative DTI-ALPS medial loading (−0.227); ReHo and CSF volumes contributed minimally (all |loading| < 0.08). PC1 thus indexes the joint extracellular-water and white-matter microstructural profile, and although it correlates substantially with global mean free water (r = +0.67, RZ = 0.44), it captures MDt and FAt variance not carried by free water alone.

**Figure 4.**
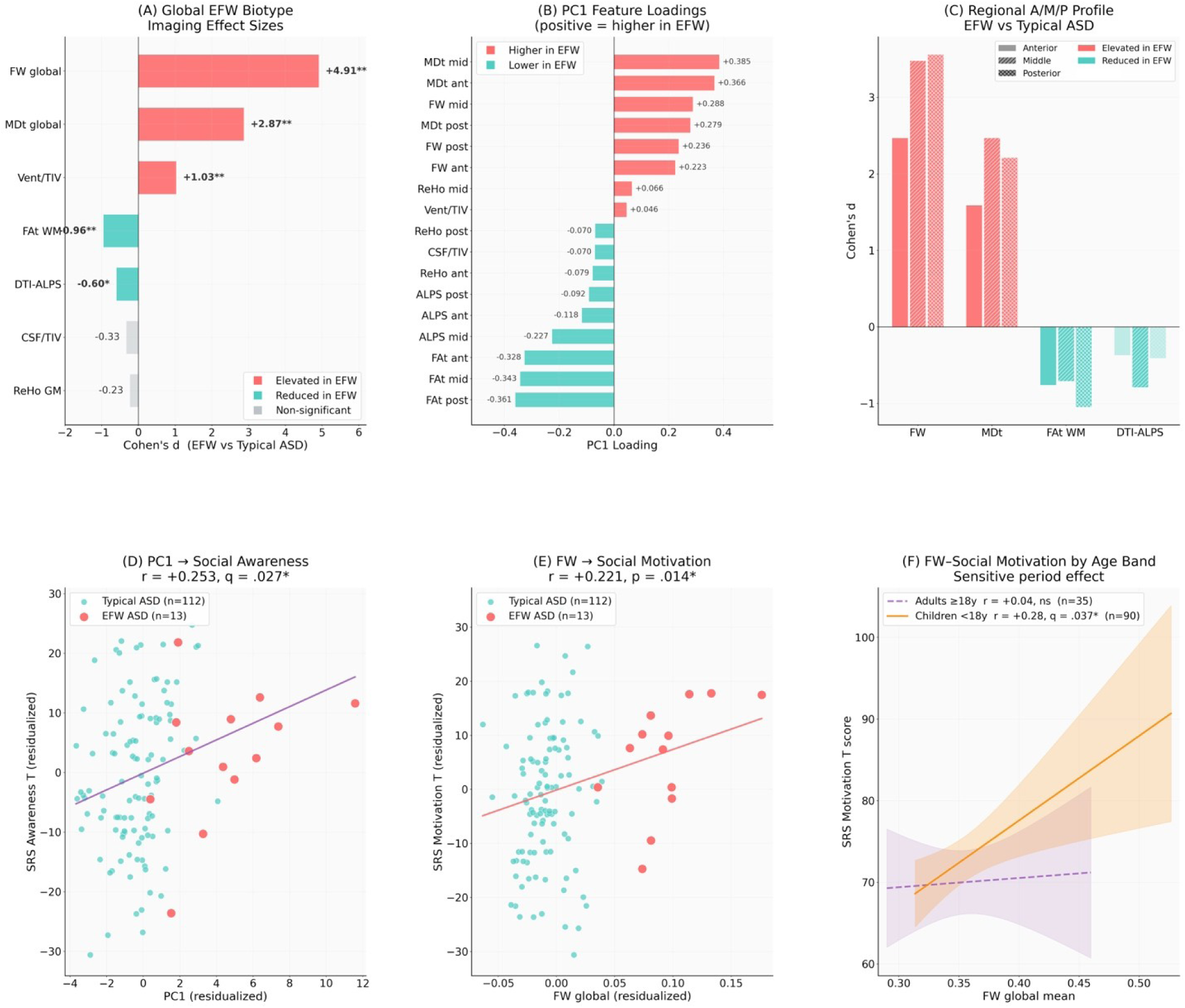
EFW biotype imaging characteristics and clinical associations. (A) Global effect sizes (partial Cohen’s d; ANCOVA covarying age and sex, plus TIV for CSF volumes) for the EFW versus non-EFW biotype contrast: FW elevation dominates (d = +4.91), followed by MDt (d = +2.87), Vent/TIV (d = +1.03), FAt WM reduction (d = −0.96), and DTI-ALPS reduction (d = −0.60). Salmon = elevated in EFW, teal = reduced in EFW, grey = non-significant. Because the biotypes were defined by clustering on these features, these effect sizes describe the cluster-defining contrast rather than independent tests; comparisons against typically developing controls are reported in the text. (B) PC1 feature loadings from the 17-feature regional PCA (positive = higher in the EFW biotype): MDt loads most positively and FAt WM most negatively (full loadings in Additional file 1: Table S10). (C) Regional anterior/medial/posterior effect-size profiles for FW, MDt, FAt WM, and DTI-ALPS: DTI-ALPS shows a medial-predominant reduction (d = −0.79**, p = .006), while FW and MDt are uniformly elevated across positions. (D) PC1 associates most strongly with SRS social awareness (partial r = +0.253, q = .026*); social cognition and social communication were also significant after FDR correction (Additional file 1: Table S11). (E) FW global mean associates with SRS social motivation across ASD (n = 125; partial r = +0.221, p = .014*). (F) The FW–social motivation association is significant in children and adolescents (< 18 years: r = +0.29, q = .037*; solid orange line, 95% CI shaded) but null in adults (≥ 18 years: r = +0.04, ns; dashed purple), despite nearly identical EFW free-water levels across age bands (children M = 0.446, adults M = 0.444). All partial correlations are residualized for age and sex. CI = 95% confidence interval; *p < .05; **p < .01.

After FDR correction within the SRS subscale family (n = 125, age and sex covaried), PC1 was positively associated with three of five subscales: Awareness (partial r = +0.253, q = .026; Figure 4D), Cognition (r = +0.201, q = .046) and Communication (r = +0.199, q = .046). Motivation (r = +0.125), Mannerisms (r = +0.083), ADOS-G total (r = +0.062) and IQ (r = +0.029) were all non-significant (full family in Additional file 1: Table S11). PC1 therefore relates to the social-perceptual and communicative dimensions of the SRS, most strongly social awareness, but not to diagnostic severity or general cognitive ability.

When PC1 and global mean free water were entered as simultaneous co-predictors (n = 125, age and sex covaried; BH FDR within family), the two dissociated. PC1 uniquely associated with SRS Awareness (β = +4.68, p = .003, q = .017; free water non-significant, p = .165), whereas free water showed unique associations with Motivation (β = +3.39, p = .037) and Mannerisms (β = +3.50, p = .080) for which PC1 was null, though neither free-water association survived FDR (both q > .22). No other subscale reached significance for either predictor (all p > .17), and both were null for ADOS-G (n = 88) and IQ (all q > .32; Additional file 1: Table S9). PC1 and free water may therefore relate to different social dimensions, social awareness versus social motivation, though this dissociation is preliminary.

### Age-Stratified FW-SRS Associations

FW-Motivation was the sole significant association in children and adolescents (r = +0.287, p = .006, q = .037; n = 90), with EFW FW levels nearly identical across age bands (children M = 0.446, adults M = 0.444), suggesting the developmental difference reflects phenotypic expression rather than imaging marker levels (Figure 4F). All remaining SRS subscales, ADOS-G, and IQ were null in both age bands (all q > .17).

## Discussion

Unsupervised clustering of multimodal MRI identified a neurobiologically distinct elevated free-water biotype within ASD. Although free water is reliably elevated in ASD at the group level [24–27], our results suggest this reflects a subgroup of roughly 10% rather than a uniform effect, reframing whole-group differences as a subset effect. This biotype is defined by fluid and tissue diffusion measures rather than the cortical morphometry or functional connectivity used thus far for ASD biotyping [2,11–14]. It extends to ASD a diffusion-MRI subtyping approach previously applied in Parkinson’s and Alzheimer’s disease [16,69], and adds perivascular diffusivity (DTI-ALPS), which to our knowledge has not previously been used as a clustering feature for stratification [70–74]. Within the EFW cluster, elevated free water, reduced anisotropy, reduced DTI-ALPS and enlarged CSF spaces together suggest a common alteration of the periventricular neurofluid and white matter environment. That these measures move together and recover across two feature sets suggests a coherent biological story rather than the single-metric findings typical in ASD research. Importantly, this signature related within ASD to social-perceptual and communicative symptoms rather than diagnostic severity or IQ. This matters because many neuroimaging measures, from atrophy to white matter differences, correlate with severity across nearly all disorders [75], a nonspecific pattern that says little about any one condition. A marker that instead relates specifically to social function suggests the underlying neurofluid biology engages the core social domain of ASD, unlike other disorders where such measures relate mainly to general cognition or disability [76–78].

### Interpreting the free-water signal and its elevation in ASD

Free water has been validated as a marker of neuroinflammation against postmortem histology, showing signatures of microglial and astrocyte activation confirmed by immunohistochemistry [21], and against *in vivo* markers of neuroinflammation and axonal injury including PET and plasma neurofilament light [79,80]. Prior evidence in ASD of immune dysregulation and microglial activation [79,80] suggests the free-water elevation in this biotype may reflect a comparable neuroinflammatory process. Supporting an extracellular-fluid interpretation, the mean diffusivity of tissue was the second-largest contributor to the EFW biotype, and free water and mean diffusivity, are typically elevated together [20] and in ASD [81,82]. Their co-elevation points more specifically to increased extracellular and interstitial water than either measure alone [83]. Free water is not specific to inflammation, however; it also rises with edema, extracellular expansion and tissue loss, so this interpretation [20], while consistent with the ASD immune literature, remains one of several and requires direct confirmation.

### Regional specificity of the reduction in DTI-ALPS

The selective reduction of the medial ALPS (d = −0.79, p = .006), localizes the effect to the lateral ventricular body, the territory of the deep medullary veins and recognized site of clearance and drainage vulnerability [48]. In the aging brain, venous collagenosis of these veins, confined to the periventricular white matter and sparing more superficial tracts, is a leading cause of periventricular white matter injury, producing extracellular fluid accumulation through raised venous pressure and impaired perivenous clearance [84–86]. Although this venous pathology is characterized mainly in older adults, the same periventricular deep medullary vein territory is selectively vulnerable in the developing brain, where impaired medullary venous drainage produces periventricular injury in preterm infants [87], and where markers of perivascular clearance, including enlarged perivascular spaces, are already reported in ASD and other neurodevelopmental conditions [87–89]. The developmental mechanism need not be the venous collagenosis seen in aging; rather, this location may represent a shared periventricular clearance bottleneck expressed through different pathologies across the lifespan. Whereas FW and MDt were elevated across all positions, the ALPS reduction was regionally restricted, suggesting that the diffusion environment at this vulnerable site is not simply a reflection of global microstructural abnormalities.

### EFW biotype and clinical expression

Within ASD, the EFW biotype’s imaging metrics related to social functioning rather than symptom severity or IQ. This is notable because the biotype’s FW and MDt increase and FAt decrease were global, suggesting social functioning may relate more to a widespread microstructural process which these metrics detect rather than changes in a specialized region. Alternatively, a global metric may be dominated by the most affected areas, which in ASD are disproportionately those serving social function [82], so a social relationship could emerge even from a uniform signal.

Social function, however, is not a single construct, and when PC1 and FW were entered together, they dissociated: PC1 uniquely related to social awareness while FW was uniquely associated with social motivation in adolescents and children. Notably, the pure extracellular-fluid marker related to social motivation in children/adolescents, which the social motivation theory frames as developmentally primary [90], while the broader microstructural and neurofluid signature related to social awareness in the whole ASD cohort, a later-developed perceptual function. This raises the possibility that the two measures map onto different levels of social processing.

The finding that the FW and social-motivation relationship was present in children and adolescents but not detectable in adults despite comparable FW levels across age bands may indicate that FW here is not merely an aging marker but that fluid elevation relates to social motivation only during the developmental window when white-matter myelination and perivascular clearance are most vulnerable, consistent with evidence that social experience and myelination are coupled during a critical period [91–93], though longitudinal data would be required to establish this.

## Limitations

Several limitations should be noted. The cross-sectional design precludes inference about trajectory, and causal relationships between the imaging findings and the clinical associations reported here cannot be established. The broad age range of the sample (5 to 64 years) is a further limitation, since FW elevation occurs physiologically at both ends of the age distribution; although the sensitivity analysis (age and sex included as clustering features at 0.5 weight, ARI = 1.000) indicates that the EFW biotype is not a simple age artifact, its mechanistic interpretation is likely to differ for children compared with older adults within the cluster. Clinical data completeness also varied across sites (SRS 84%, ADOS 59%, BDI-II 29%), which, together with the small size of the EFW biotype (n = 14 to 15), limited power for the imaging-clinical analyses.

Because the biotype was defined by clustering on the same imaging features subsequently evaluated with effect sizes, these within-biotype effect sizes are a consequence of cluster formation rather than independent tests, and we are reporting them as descriptive rather than inferential. The comparison against typically developing controls, who were not included in the clustering, is not subject to this circularity. Replication in independent samples, with longitudinal designs and complete clinical data, is therefore needed.

Several imaging-clinical associations were modest in size (partial r of approximately 0.2 to 0.3). Although they survived false discovery rate correction within each outcome family, most would not meet the more stringent threshold of p < .005 that has been proposed for biological psychiatry, and in the whole ASD sample the free-water association with social motivation did not survive correction. These associations should therefore be regarded as preliminary. The sample was also predominantly male, reflecting the composition of ABIDE II, which limits generalization to females with ASD. Finally, the primary findings could not be replicated in an independent cohort within this study. ABIDE I does not include diffusion MRI, and the present analysis already includes every ABIDE II site that provided diffusion data; confirmation in an independent sample with diffusion, resting-state fMRI and dimensional social measures is the most important next step.

## Conclusions

Unsupervised clustering of multimodal MRI identified an elevated free-water biotype in approximately 10% of individuals with ASD, defined by convergent elevation of extracellular free water, reduced tissue anisotropy, reduced periventricular DTI-ALPS and enlarged ventricles. The biotype departed from typically developing norms whereas the remaining participants with ASD did not, suggesting that group-level free-water differences in autism may reflect a subgroup rather than a uniform effect. Within ASD, the imaging signature was related to social-perceptual and communicative symptoms rather than to diagnostic severity or IQ, and free water was related to social motivation in children and adolescents. If replicated in independent and longitudinal samples, this signature may prove useful as an objective marker for stratifying individuals with ASD in mechanistic studies and in trials targeting neuroinflammatory or neurofluid-related pathways.

## Declarations

### Ethics approval and consent to participate

This study was a secondary analysis of de-identified data from the Autism Brain Imaging Data Exchange II (ABIDE II). Each contributing site collected data under protocols approved by its local institutional review board, with informed consent obtained from participants or their legal guardians, and shared the data in de-identified form. Because only publicly available, de-identified data were analyzed, no additional ethics approval was required.

### Consent for publication

Not applicable.

### Availability of data and materials

The data analysed during the current study are publicly available from the ABIDE II repository of the International Neuroimaging Data-sharing Initiative [50].

### Authors’ contributions

FM conceptualized the study, developed the methodology, performed the formal analysis and visualization, and wrote the original draft. KH curated the data. SN developed software and curated the data. SM and DZ contributed to study conceptualization, supervised the work, and provided resources and funding. All authors reviewed and edited the manuscript, and read and approved the final manuscript.

## Acknowledgements

ABIDE-II data were provided by the Autism Brain Imaging Data Exchange II consortium. Primary support for the work by Adriana Di Martino and her team was provided by the National Institute of Mental Health (NIMH 5R21MH107045). Primary support for the work by Michael P. Milham and his team was provided by the National Institute of Mental Health (NIMH 5R21MH107045) and the Nathan S. Kline Institute for Psychiatric Research. Additional support was provided by gifts from Joseph P. Healey, Phyllis Green, and Randolph Cowen to the Child Mind Institute.

## Additional files

**Additional file 1 (.docx):** Supplementary Information. Supplementary methods (data selection and quality control, regional parcellation, DTI-ALPS, free water imaging, rs-fMRI and harmonization), supplementary results, Supplementary Tables S1 to S11 and Supplementary Figures S1 and S2.

## Abbreviations

aALPS: anterior DTI-ALPS index
ABIDE: Autism Brain Imaging Data Exchange
ADI-R: Autism Diagnostic Interview-Revised
ADOS-G: Autism Diagnostic Observation Schedule-Generic
ADt: tissue (free-water-corrected) axial diffusivity
AFNI: Analysis of Functional NeuroImages
ALFF: amplitude of low-frequency fluctuations
ANCOVA: analysis of covariance
ARI: adjusted Rand index
ASD: autism spectrum disorder
BDI-II: Beck Depression Inventory-II
BH: Benjamini-Hochberg
BIC: Bayesian information criterion
BNI: Barrow Neurological Institute
BOLD: blood oxygen level dependent
CI: confidence interval
CSF: cerebrospinal fluid
DTI: diffusion tensor imaging
DTI-ALPS: diffusion tensor image analysis along the perivascular space
EFW: elevated free water
FA: fractional anisotropy
fALFF: fractional amplitude of low-frequency fluctuations
FAt: tissue (free-water-corrected) fractional anisotropy
FDR: false discovery rate
FIQ: full-scale IQ
fMRI: functional magnetic resonance imaging
FSL: FMRIB Software Library
FW: free water
FWE: free water elimination
FWI: free water imaging
gALPS: global DTI-ALPS index
GM: gray matter
GMM: Gaussian mixture model
IDDRC: Intellectual and Developmental Disabilities Research Center
IP: Institut Pasteur and Robert Debré Hospital
IQ: intelligence quotient
JHU: Johns Hopkins University
mALPS: medial DTI-ALPS index
MD: mean diffusivity
MDt: tissue (free-water-corrected) mean diffusivity
MNI: Montreal Neurological Institute
MRI: magnetic resonance imaging
NYU: New York University
OLS: ordinary least squares
pALPS: posterior DTI-ALPS index
PC: principal component
PCA: principal component analysis
PET: positron emission tomography
PIQ: performance IQ
RDt: tissue (free-water-corrected) radial diffusivity
ReHo: regional homogeneity
ROI: region of interest
SD: standard deviation
SDSU: San Diego State University
SRS: Social Responsiveness Scale
TCD: Trinity College Dublin
TD: typically developing
TIV: total intracranial volume
VIQ: verbal IQ
WM: white matter

## References

1. Lombardo M V., Lai MC, Baron-Cohen S. Big data approaches to decomposing heterogeneity across the autism spectrum. Mol Psychiatry. 2019;24:1435–1450.

2. Hong SJ, Vogelstein JT, Gozzi A, Bernhardt BC, Yeo BTT, Milham MP, et al. Toward Neurosubtypes in Autism. Biol Psychiatry. 2020;88:111–128.

3. Parellada M, Andreu-Bernabeu Á, Burdeus M, Cáceres ASJ, Urbiola E, Carpenter LL, et al. In Search of Biomarkers to Guide Interventions in Autism Spectrum Disorder: A Systematic Review. Am J Psychiatry. 2023;180:23–40.

4. Feczko E, Miranda-Dominguez O, Marr M, Graham AM, Nigg JT, Fair DA. The Heterogeneity Problem: Approaches to Identify Psychiatric Subtypes. Trends Cogn Sci. 2019;23:584–601.

5. Drysdale AT, Grosenick L, Downar J, Dunlop K, Mansouri F, Meng Y, et al. Resting-state connectivity biomarkers define neurophysiological subtypes of depression. Nat Med. 2017;23:28–38.

6. Tamminga CA, Pearlson G, Keshavan M, Sweeney J, Clementz B, Thaker G. Bipolar and schizophrenia network for intermediate phenotypes: Outcomes across the psychosis continuum. Schizophr Bull. 2014;40 Suppl 2:S131–S137.

7. Tamminga CA, Ivleva EI. Using Brain-Based Phenotyping to Improve Discovery in Psychiatry. JAMA Psychiatry. 2018;75:1103–1104.

8. Eshaghi A, Young AL, Wijeratne PA, Prados F, Arnold DL, Narayanan S, et al. Identifying multiple sclerosis subtypes using unsupervised machine learning and MRI data. Nat Commun. 2021;12:2078.

9. Young AL, Marinescu R V., Oxtoby NP, Bocchetta M, Yong K, Firth NC, et al. Uncovering the heterogeneity and temporal complexity of neurodegenerative diseases with Subtype and Stage Inference. Nat Commun. 2018;9:4273.

10. Ferreira D, Nordberg A, Westman E. Biological subtypes of Alzheimer disease: A systematic review and meta-analysis. Neurology. 2020;94:436–448.

11. Urchs SGW, Tam A, Orban P, Moreau C, Benhajali Y, Nguyen HD, et al. Functional connectivity subtypes associate robustly with ASD diagnosis. Elife. 2022;11:e56257.

12. Chen H, Uddin LQ, Guo X, Wang J, Wang R, Wang X, et al. Parsing brain structural heterogeneity in males with autism spectrum disorder reveals distinct clinical subtypes. Hum Brain Mapp. 2019;40:628–637.

13. Pagani M, Zerbi V, Gini S, Alvino FG, Banerjee A, Barberis A, et al. Autism subtypes identified using cross-species functional connectivity analyses. Nat Neurosci. 2026;29:1476–1487.

14. Buch AM, Vértes PE, Seidlitz J, Kim SH, Grosenick L, Liston C. Molecular and network-level mechanisms explaining individual differences in autism spectrum disorder. Nat Neurosci. 2023;26:650–663.

15. Burciu RG, Ofori E, Archer DB, Wu SS, Pasternak O, McFarland NR, et al. Progression marker of Parkinson’s disease: A 4-year multi-site imaging study. Brain. 2017;140:2183–2192.

16. Vijayakumari AA, Sakaie KE, Teixeira-Dos-Santos D, Fernandez HH, Walter BL. Data-driven neurobiological subtyping of Parkinson’s disease using diffusion MRI-derived isotropic diffusion. Neuroradiology. 2026;68:e-pub ahead of print.

17. Assaf Y. Imaging laminar structures in the gray matter with diffusion MRI. Neuroimage. 2019;197:677–688.

18. Maillard P, Fletcher E, Singh B, Martinez O, Johnson DK, Olichney JM, et al. Cerebral white matter free water: A sensitive biomarker of cognition and function. Neurology. 2019;92:e2221– e2231.

19. Cai Y, Zhang Y, Leng S, Ma Y, Jiang Q, Wen Q, et al. The relationship between inflammation, impaired glymphatic system, and neurodegenerative disorders: A vicious cycle. Neurobiol Dis. 2024;192:106426.

20. Pasternak O, Sochen N, Gur Y, Intrator N, Assaf Y. Free water elimination and mapping from diffusion MRI. Magn Reson Med. 2009;62:717–730.

21. Garcia-Hernandez R, Cerdá AC, Carpena AT, Drakesmith M, Koller K, Jones DK, et al. Mapping microglia and astrocyte activation in vivo using diffusion MRI. Sci Adv. 2022;8:eabq2923.

22. Febo M, Perez PD, Ceballos-Diaz C, Colon-Perez LM, Zeng H, Ofori E, et al. Diffusion magnetic resonance imaging-derived free water detects neurodegenerative pattern induced by interferon-γ. Brain Struct Funct. 2020;225:427–439.

23. Di Biase MA, Katabi G, Piontkewitz Y, Cetin-Karayumak S, Weiner I, Pasternak O. Increased extracellular free-water in adult male rats following in utero exposure to maternal immune activation. Brain Behav Immun. 2020;83:283–287.

24. Walsh MJM, Ofori E, Pagni BA, Chen K, Sullivan G, Braden BB. Preliminary findings of accelerated visual memory decline and baseline brain correlates in middle-age and older adults with autism: The case for hippocampal free-water. Front Aging Neurosci. 2022;14:1029166.

25. Zhang Y, Ofori E, Chen K, Harker SA, Velez M, Gallegos S, et al. Association of glymphatic dysfunction, free water, white matter integrity and long-term memory performance in aging autistic adults. Mol Autism. 2026;17:10.

26. Wilkes BJ, Archer DB, Farmer AL, Bass C, Korah H, Vaillancourt DE, et al. Cortico-basal ganglia white matter microstructure is linked to restricted repetitive behavior in autism spectrum disorder. Mol Autism. 2024;15:6.

27. Surgent O, Riaz A, Ausderau KK, Adluru N, Kirk GR, Guerrero-Gonzalez J, et al. Brainstem white matter microstructure is associated with hyporesponsiveness and overall sensory features in autistic children. Mol Autism. 2022;13:48.

28. Shen MD, Kim SH, McKinstry RC, Gu H, Hazlett HC, Nordahl CW, et al. Increased Extra-axial Cerebrospinal Fluid in High-Risk Infants Who Later Develop Autism. Biol Psychiatry. 2017;82:186–193.

29. Peterson M, Prigge MBD, Bigler ED, Zielinski B, King JB, Lange N, et al. Evidence for normal extra-axial cerebrospinal fluid volume in autistic males from middle childhood to adulthood. Neuroimage. 2021;240:118387.

30. Shen MD. Cerebrospinal fluid and the early brain development of autism. J Neurodev Disord. 2018;10:39.

31. Iliff JJ, Wang M, Liao Y, Plogg BA, Peng W, Gundersen GA, et al. A paravascular pathway facilitates CSF flow through the brain parenchyma and the clearance of interstitial solutes, including amyloid β. Sci Transl Med. 2012;4:147ra111.

32. Rasmussen MK, Mestre H, Nedergaard M. Fluid transport in the brain. Physiol Rev. 2022;102:1025–1151.

33. Moody DM, Bell MA, Challa VR. Features of the cerebral vascular pattern that predict vulnerability to perfusion or oxygenation deficiency: An anatomic study. AJNR Am J Neuroradiol. 1990;11:431–439.

34. Taoka T, Ito R, Nakamichi R, Nakane T, Naganawa S. Diffusion tensor image analysis along the perivascular space (DTI-ALPS): redefining its interpretation and role. Jpn J Radiol. 2026:e-pub ahead of print.

35. Taoka T, Masutani Y, Kawai H, Nakane T, Matsuoka K, Yasuno F, et al. Evaluation of glymphatic system activity with the diffusion MR technique: diffusion tensor image analysis along the perivascular space (DTI-ALPS) in Alzheimer’s disease cases. Jpn J Radiol. 2017;35:172–178.

36. Wang M, He K, Zhang L, Xu D, Li X, Wang L, et al. Assessment of glymphatic function and white matter integrity in children with autism using multi-parametric MRI and machine learning. Eur Radiol. 2025;35:3653–3663.

37. Zhao S, Zhang M, Luo T, Li L, Jiang Y, Situ M, et al. Glymphatic system dysfunction in children with autism spectrum disorder as evidenced by the diffusion tensor imaging along perivascular spaces index. Front Psychiatry. 2025;16:1701816.

38. Li X, Ruan C, Zibrila AI, Musa M, Wu Y, Zhang Z, et al. Children with autism spectrum disorder present glymphatic system dysfunction evidenced by diffusion tensor imaging along the perivascular space. Medicine (Baltimore). 2022;101:e32061.

39. Costa T, Manuello J, Premi E, Mattioli I, Lasagna L, Lahoz CB, et al. Evaluating the robustness of DTI-ALPS in clinical context: a meta-analytic parallel on Alzheimer’s and Parkinson’s diseases. Sci Rep. 2024;14:26381.

40. Zhang F, Li Q, Wang M, Zhang Y, Chen J, Shang X. A systematic review and meta-analysis on glymphatic flow dysfunction in depression:Evidence from the DTI-ALPS index. J Affect Disord. 2026;410:121960.

41. Di Martino A, Yan CG, Li Q, Denio E, Castellanos FX, Alaerts K, et al. The autism brain imaging data exchange: Towards a large-scale evaluation of the intrinsic brain architecture in autism. Mol Psychiatry. 2014;19:659–667.

42. Fortin J-P, Cullen N, Sheline YI, Taylor WD, Aselcioglu I, Cook PA, et al. Harmonization of cortical thickness measurements across scanners and sites. Neuroimage. 2018;167:104–120.

43. Fortin JP, Parker D, Tunç B, Watanabe T, Elliott MA, Ruparel K, et al. Harmonization of multi-site diffusion tensor imaging data. Neuroimage. 2017;161:149–170.

44. Zang Y, Jiang T, Lu Y, He Y, Tian L. Regional homogeneity approach to fMRI data analysis. Neuroimage. 2004;22:394–400.

45. Sui J, Zhi D, Calhoun VD. Data-driven multimodal fusion: Approaches and applications in psychiatric research. Psychoradiology. 2023;3:kkad026.

46. Calhoun VD, Sui J. Multimodal Fusion of Brain Imaging Data: A Key to Finding the Missing Link(s) in Complex Mental Illness. Biol Psychiatry Cogn Neurosci Neuroimaging. 2016;1:230– 244.

47. Clementz BA, Sweeney JA, Hamm JP, Ivleva EI, Ethridge LE, Pearlson GD, et al. Identification of distinct psychosis biotypes using brain-based biomarkers. Am J Psychiatry. 2016;173:373– 384.

48. Taoka T, Fukusumi A, Miyasaka T, Kawai H, Nakane T, Kichikawa K, et al. Structure of the medullary veins of the cerebral hemisphere and related disorders. Radiographics. 2017;37:281– 297.

49. Zhang R, Zhou Y, Yan S, Zhong G, Liu C, Jiaerken Y, et al. A brain region-based deep medullary veins visual score on susceptibility weighted imaging. Front Aging Neurosci. 2017;9:269.

50. Di Martino A, O’Connor D, Chen B, Alaerts K, Anderson JS, Assaf M, et al. Enhancing studies of the connectome in autism using the autism brain imaging data exchange II. Sci Data. 2017;4:170010.

51. Lord C, Risi S, Lambrecht L, Cook EH, Leventhal BL, Dilavore PC, et al. The Autism Diagnostic Observation Schedule-Generic: A standard measure of social and communication deficits associated with the spectrum of autism. J Autism Dev Disord. 2000;30:205–223.

52. Lord C, Rutter M, Le Couteur A. Autism Diagnostic Interview-Revised: A revised version of a diagnostic interview for caregivers of individuals with possible pervasive developmental disorders. J Autism Dev Disord. 1994;24:659–685.

53. Constantino JN, Gruber CP. Social Responsiveness Scale, Second Edition (SRS-2). Los Angeles, CA: Western Psychological Services; 2012.

54. Kaufman AS, Kaufman NL. Kaufman Brief Intelligence Test, Second Edition. Encyclopedia of Special Education, Bloomington, MN: NCS Pearson; 2004.

55. Wechsler D. Manual for the Wechsler abbreviated intelligence scale (WASI). San Antonio, TX: Psychological Corporation; 1999.

56. Kaufman AS, Flanagan DP, Alfonso VC, Mascolo JT. Test Review: Wechsler Intelligence Scale for Children, Fourth Edition (WISC-IV). J Psychoeduc Assess. 2006;24:278–295.

57. Smith SM, Jenkinson M, Woolrich MW, Beckmann CF, Behrens TEJ, Johansen-Berg H, et al. Advances in functional and structural MR image analysis and implementation as FSL. Neuroimage. 2004;23:S208–S219.

58. Zhang Y, Brady M, Smith S. Segmentation of brain MR images through a hidden Markov random field model and the expectation-maximization algorithm. IEEE Trans Med Imaging. 2001;20:45–57.

59. Tournier J-D, Smith R, Raffelt D, Tabbara R, Dhollander T, Pietsch M, et al. MRtrix3: a fast, flexible and open software framework for medical image processing and visualisation. Neuroimage. 2019;202:116137.

60. Jenkinson M, Beckmann CF, Behrens TEJ, Woolrich MW, Smith SM. FSL. Neuroimage. 2012;62:782–790.

61. Schilling KG, Blaber J, Huo Y, Newton A, Hansen C, Nath V, et al. Synthesized b0 for diffusion distortion correction (Synb0-DisCo). Magn Reson Imaging. 2019;64:62–70.

62. Fonov V, Evans AC, Botteron K, Almli CR, McKinstry RC, Collins DL. Unbiased average age-appropriate atlases for pediatric studies. Neuroimage. 2011;54:313–327.

63. Jiang D, Liu L, Kong Y, Chen Z, Rosa-Neto P, Chen K, et al. Regional Glymphatic Abnormality in Behavioral Variant Frontotemporal Dementia. Ann Neurol. 2023;94:442–456.

64. Golub M, Neto Henriques R, Gouveia Nunes R. Free-water DTI estimates from single b-value data might seem plausible but must be interpreted with care. Magn Reson Med. 2021;85:2537– 2551.

65. Cox RW. AFNI: Software for analysis and visualization of functional magnetic resonance neuroimages. Comput Biomed Res. 1996;29:162–173.

66. Friston KJ, Williams S, Howard R, Frackowiak RSJ, Turner R. Movement-related effects in fMRI time-series. Magn Reson Med. 1996;35:346–355.

67. Benjamini Y, Hochberg Y. Controlling the false discovery rate: a practical and powerful approach to multiple testing. J R Stat Soc Series B Stat Methodol. 1995;57:289–300.

68. Carreira Figueiredo I, Borgan F, Pasternak O, Turkheimer FE, Howes OD. White-matter free-water diffusion MRI in schizophrenia: a systematic review and meta-analysis. Neuropsychopharmacology. 2022;47:1413–1420.

69. Habes M, Grothe MJ, Tunc B, McMillan C, Wolk DA, Davatzikos C. Disentangling Heterogeneity in Alzheimer’s Disease and Related Dementias Using Data-Driven Methods. Biol Psychiatry. 2020;88:70–82.

70. Kim BH, Shin D, Kang SH, Jang H, Yun J, Park YH, et al. Mapping Alzheimer’s disease pathology using free water through integrated analysis of plasma biomarkers, microstructural DTI metrics, and macrostructural MRI measures. Sci Rep. 2025;15:30702.

71. Hoy AR, Ly M, Carlsson CM, Okonkwo OC, Zetterberg H, Blennow K, et al. Microstructural white matter alterations in preclinical Alzheimer’s disease detected using free water elimination diffusion tensor imaging. PLoS One. 2017;12:e0173982.

72. Bergamino M, McElvogue MM, Stokes AM. Distinguishing Early from Late Mild Cognitive Impairment Using Magnetic Resonance Free-Water Diffusion Tensor Imaging. NeuroSci. 2025;6:8.

73. Zhou G, Ren J, Rong D, Zhou H, Ning H, Wang H, et al. Monitoring Substantia Nigra Degeneration Using Free Water Imaging across Prodromal and Clinical Parkinson’s Disease. Mov Disord. 2023;38:774–782.

74. Guttuso T, Bergsland N, Hagemeier J, Lichter DG, Pasternak O, Zivadinov R. Substantia nigra free water increases longitudinally in Parkinson disease. AJNR Am J Neuroradiol. 2018;39:479– 484.

75. Romer AL, Elliott ML, Knodt AR, Sison ML, Ireland D, Houts R, et al. Pervasively thinner neocortex as a transdiagnostic feature of general psychopathology. Am J Psychiatry. 2021;178:174–182.

76. Zhang X, Hatoum L, Ying J, Huang C. Assessing glymphatic-associated fluid dynamics in psychiatric disorders: evidence from neuroimaging - a review. Psychoradiology. 2025;5:kkaf031.

77. Abdolizadeh A, Torres-Carmona E, Kambari Y, Amaev A, Song J, Ueno F, et al. Evaluation of the Glymphatic System in Schizophrenia Spectrum Disorder Using Proton Magnetic Resonance Spectroscopy Measurement of Brain Macromolecule and Diffusion Tensor Image Analysis Along the Perivascular Space Index. Schizophr Bull. 2024;50:1396–1410.

78. Fu Q, Qiu Y, Hu Y, Wang Y, Wang Y, Hu W, et al. Free Water-Corrected Fractional Anisotropy in Normal-Appearing White Matter as a Potential Neuroimaging Biomarker for Attention and Executive Function Impairment in Cerebral Small Vessel Disease. CNS Neurosci Ther. 2025;31:e70475.

79. Vargas DL, Nascimbene C, Krishnan C, Zimmerman AW, Pardo CA. Neuroglial activation and neuroinflammation in the brain of patients with autism. Ann Neurol. 2005;57:67–81.

80. Morgan JT, Chana G, Pardo CA, Achim C, Semendeferi K, Buckwalter J, et al. Microglial activation and increased microglial density observed in the dorsolateral prefrontal cortex in autism. Biol Psychiatry. 2010;68:368–376.

81. Andica C, Kamagata K, Kirino E, Uchida W, Irie R, Murata S, et al. Neurite orientation dispersion and density imaging reveals white matter microstructural alterations in adults with autism. Mol Autism. 2021;12:48.

82. Aoki Y, Abe O, Nippashi Y, Yamasue H. Comparison of white matter integrity between autism spectrum disorder subjects and typically developing individuals: A meta-analysis of diffusion tensor imaging tractography studies. Mol Autism. 2013;4:25.

83. Hu S, Exner C, Sienel RI, When AC, Seker FB, Magdane Boldoczki F, et al. Characterization of Vasogenic and Cytotoxic Brain Edema Formation After Experimental Traumatic Brain Injury by Free Water Diffusion Magnetic Resonance Imaging. J Neurotrauma. 2024;41:393–406.

84. Gao F, Ramirez J, Holmes MF, Keith J, Masellis M, Swartz RH, et al. Periventricular white matter hyperintensities co-localize with deep intramedullary venules, sparing compact fiber tracts and can increase or decrease over time in Alzheimer’s disease. Alzheimers Dement. 2025;41:e099476.

85. Moody DM, Brown WR, Challa VR, Anderson RL. Periventricular venous collagenosis: Association with leukoaraiosis. Radiology. 1995;194.

86. Keith J, Gao FQ, Noor R, Kiss A, Balasubramaniam G, Au K, et al. Collagenosis of the deep medullary veins: An underrecognized pathologic correlate of white matter hyperintensities and periventricular infarction? J Neuropathol Exp Neurol. 2017;76:299–312.

87. Chen D, Sun J, Li Q, Bai W, Mao J. An Important Finding of White Matter Injury in Late Preterm Infant: Deep Medullary Vein Involvement. Front Pediatr. 2020;8:597567.

88. Garic D, McKinstry RC, Rutsohn J, Slomowitz R, Wolff J, MacIntyre LC, et al. Enlarged Perivascular Spaces in Infancy and Autism Diagnosis, Cerebrospinal Fluid Volume, and Later Sleep Problems. JAMA Netw Open. 2023;6:e2348341.

89. Chen Y, Wang M, Su S, Dai Y, Zou M, Lin L, et al. Assessment of the glymphatic function in children with attention-deficit/hyperactivity disorder. Eur Radiol. 2024;34:1444–1452.

90. Chevallier C, Kohls G, Troiani V, Brodkin ES, Schultz RT. The social motivation theory of autism. Trends Cogn Sci. 2012;16:231–239.

91. Schneider N, Greenstreet E, Deoni SCL. Connecting inside out: Development of the social brain in infants and toddlers with a focus on myelination as a marker of brain maturation. Child Dev. 2022;93:359–371.

92. Makinodan M, Rosen KM, Ito S, Corfas G. A critical period for social experience-dependent oligodendrocyte maturation and myelination. Science (1979). 2012;337:1357–1360.

93. Supekar K, Kochalka J, Schaer M, Wakeman H, Qin S, Padmanabhan A, et al. Deficits in mesolimbic reward pathway underlie social interaction impairments in children with autism. Brain. 2018;141:2795–2805.

