## Supplemental Material for "Multimodal MRI Clustering Reveals an Elevated Free-Water Biotype in Autism with Increased Social Symptoms"

McKenna et al.

### S1. Supplementary Methods

#### S1.1 Data Selection

Data were drawn from the Autism Brain Imaging Data Exchange II (ABIDE II) [1], an open access, multi-site repository established to aggregate resting state functional and structural neuroimaging data across autism research sites. The present analysis included data from six contributing samples: Barrow Neurological Institute (BNI), New York University Langone Medical Center (NYU1 and NYU2), Institut Pasteur and Robert Debré Hospital (IP), San Diego State University (SDSU), and Trinity College Dublin (TCD). Sites were selected based on availability of diffusion weighted imaging (DWI) data.

Quality control proceeded in three stages. In the first stage, raw structural and diffusion data were screened for the presence of all required acquisition files, complete header information, and matching b value/b vector tables. Subjects with missing or corrupted files were excluded prior to preprocessing. In the second stage, structural T1 weighted images were visually inspected for excessive motion artifact, ringing, ghosting, intensity inhomogeneity exceeding the range correctable by N4 bias field correction, and gross anatomical abnormalities. Diffusion data were screened for slice dropout, excessive in volume motion using framewise displacement metrics derived during eddy current and motion correction (FSL eddy)[2] , and signal to noise ratio (SNR) within the corpus callosum. Subjects with framewise displacement greater than 2 mm in more than 20% of volumes, or SNR values more than 3 standard deviations below the site mean, were flagged for exclusion. In the third stage, processed diffusion outputs were reviewed for registration accuracy to MNI space, the presence of artifactual signal in the ventricular system, and successful tensor fitting across the white matter skeleton. Final quality control disposition was then determined.

Following the full quality control pipeline, 278 participants were retained for analysis: 149 ASD (mean age = 16.5 ± 13.5 years; 128 M / 18 F; full scale IQ = 103.5 ± 17.6) and 129 TD (mean age = 21.3 ± 14.5 years; 104 M / 24 F; full scale IQ = 112.2 ± 14.4). Group level demographic and clinical characteristics are presented in Table 1 of the main text.

**S1.2 Neurocombat**

Multi-site neuroimaging data introduce systematic non-biological variance arising from differences in scanner hardware, acquisition protocols, and reconstruction software. Left unaddressed, site effects confound group comparisons and inflate clustering solutions by creating site-driven subgroups rather than neurobiologically meaningful ones. neuroCombat [3,4] was applied to remove additive and multiplicative site effects while preserving biological variance of interest. neuroCombat was applied using the empirical Bayes parametric method (eb = TRUE, parametric = TRUE), correcting for both mean (location) and variance (scale) site effects. Six ComBat batches were defined corresponding to the six ABIDE II acquisition sites included in this study: BNI, NYU1, NYU2, IP, SDSU, and TCD. Harmonization was performed on all 75 neuroimaging features simultaneously, including DTI-ALPS indices, free water (FW), tissue axial diffusivity (ADt), tissue mean diffusivity (MDt), tissue fractional anisotropy (FAt), CSF and structural volumes, and resting-state ReHo metrics (included as an exploratory functional feature). Biological covariates of interest (group: ASD/TD; age; sex) were included in the ComBat model to protect their variance from removal during harmonization. Site effects before and after harmonization are summarized in Table S1.

Site effects were quantified before and after harmonization using one-way ANOVA with site as factor, with effect size expressed as eta-squared ($\eta^2$ = SS_site / SS_total). A feature was considered to have a significant site effect if the ANOVA reached $p < .05$. Post-harmonization site effects were assessed using the same procedure on the ComBat-corrected data.

#### S1.3 DTI-ALPS Ventricle-Fitting Procedure

To account for individual differences in ventricular size, which is a particular concern in neurodevelopmental populations, projection ROIs were adjusted using a subject-specific ventricle-fitting procedure. Each subject's lateral ventricle volume was estimated from FSL's fslanat output, using FIRST subcortical segmentation labels (bilateral lateral ventricles and third/fourth ventricles) as the primary source, with fallback to FIRST individual structure files or FAST CSF partial-volume estimates (PVE > 0.3, restricted to brain interior) [5]. The subject's ventricle volume was then compared to the Harvard-Oxford atlas ventricle volume (HO labels 3, 4, 14, 15), and the atlas mask was dilated or eroded by the cube root of the volume ratio (capped at 10/5 iterations for dilation/erosion).

Projection ROIs were subsequently shifted along the mediolateral axis to track the scaled ventricle wall, with a fixed 2 mm clearance gap enforced between the ventricular wall and the nearest ROI edge, and any ROI voxels overlapping the (scaled) ventricle were removed. Association ROIs were not shifted but were similarly masked to exclude ventricular voxels. This approach avoids subject-to-MNI warping artifacts while still capturing individual anatomical variation.

ROIs were further constrained along the rostrocaudal axis to span the lateral ventricular body (Y range auto-detected from the HO atlas at Z = +26, ±10 mm medullary vein buffer [6] beyond the ventricle extent), then divided into three equal thirds yielding anterior (aDTI-ALPS), medial (mDTI-ALPS), and posterior (pDTI-ALPS) sub-indices. A complementary gradient analysis was performed by computing the ALPS index at consecutive $3\times3\times3$ mm$^3$ ROI positions along the rostrocaudal axis in 1 mm steps along the ROI, yielding a continuous anterior-to-posterior ALPS profile.

The DTI-ALPS index was computed from tensor component maps (Dxx, Dyy, Dzz) as: [(Dxx_proj + Dxx_assoc)/2] / [(Dyy_proj + Dzz_assoc)/2]. Bilateral regional indices were averaged, and global ALPS (gDTI-ALPS) was computed as the mean across all regions. The FW-corrected DTI-ALPS indices were computed analogously from the FWI-derived tensors.

**S1.4 CSF Volume Quantification**

The total CSF partial volume estimate from FSL FAST [5] was thresholded at >0.5 and decomposed into connected components using 3D labeling (scipy.ndimage.label) [7]. Components were classified as ventricular based on cluster size (>50 voxels) and proximity to the brain centroid (normalized Euclidean distance <0.25), capturing the lateral ventricles while excluding sulcal and cisternal CSF. In cases where this approach yielded insufficient ventricular voxels, a fallback method intersected CSF-thresholded voxels with a morphologically eroded brain mask, retaining only deep interior CSF. Extraventricular CSF volume was computed as total minus ventricular CSF volume. Total brain volume was computed as the sum of all tissue partial volume estimates (CSF + GM + WM). All volumes were expressed in mm$^3$ (Figure 1C of the main text).

**S1.5 Feature Selection**

Feature selection for the global and regional clustering sets was performed in two stages. First, pairwise Pearson correlations were computed among all 11 candidate global features to assess collinearity (Supplementary Table S4). Second, a drop-one analysis was conducted on the global candidate set: for each feature, it was removed from the set and k-means clustering (k = 2, 200 seeds, best silhouette selected) was re-run on z-scored, PCA-reduced features (80% variance threshold). The change in silhouette (Δ sil) relative to the full set was recorded. Features that substantially reduced silhouette when removed (Δ sil < −0.05) were prioritized for retention. Features that were added individually to the set to evaluate potential inclusion were assessed by the same criterion. The final global set (n = 7) and regional set (n = 17; positional A/M/P expansion of global diffusion features plus two CSF/TIV measures) were selected based on this combined criterion and theoretical considerations.

CSF volumes (ventricular and extraventricular) were TIV-normalized prior to inclusion. Raw ventricular volume correlated strongly with TIV (r = +0.617), which was reduced to r = +0.204 following TIV-normalization; extraventricular CSF/TIV correlation with TIV dropped from r = +0.543 to r = +0.051. TIV-normalized values were additionally winsorized at ±3 SD to remove two ASD subjects with extreme extraventricular CSF/TIV values (z-scores of 5.9).

### S2. Supplementary Analyses

#### S2.1 FWI Fit Details

For FWI estimation we used the MD-based initialization with the tissue MD prior calibrated to this cohort's deep white matter MD distribution: a conventional DTI fit was first performed, and the median MD across deep white matter voxels was computed per subject. The cross-subject median of these values (0.52 $\mu m^2/ms$; subject range: 0.50-0.56) was used as the MD reference prior for all subjects, reflecting the suppressed apparent Gaussian MD observed at b = 2500 $s/mm^2$ relative to values at lower b-values due to non-Gaussian diffusion [8,9]. Beltrami gradient descent was run for 200 iterations (learning rate = 0.0005) with the Beltrami spatial regularization term active over the first 100 iterations and disabled thereafter, following Golub et al. (https://github.com/mvgolub/FW-DTI-Beltrami) [8]. This yielded maps of free water fraction (FW) and FW-corrected fractional anisotropy (FAt), mean diffusivity (MDt), axial diffusivity (ADt), and radial diffusivity (RDt). Metrics were extracted from whole-brain and anterior/medial/posterior parcellations.

To assess whether the Beltrami FWE pipeline [8] reliably recovers free water fraction under our acquisition protocol, we performed a numerical simulation matched to the in-vivo gradient scheme. We generated 600 synthetic voxels using the exact b-value and direction table acquired for this cohort (1 b = 0 plus 32 b = 2500 s/mm$^2$ directions). Each voxel was assigned a tissue diffusion tensor with fractional anisotropy drawn uniformly from [0.1, 0.75] and mean diffusivity drawn from N(0.52, 0.05) µm$^2$/ms (matching the cohort-calibrated MD prior reported in the main Methods), with a random principal eigenvector orientation. The bi-tensor signal was constructed as a weighted sum of this tissue compartment and an isotropic free water compartment (D_iso = 3.0 × $10^{-3}$ mm$^2$/s), with the free water volume fraction drawn uniformly from [0, 0.95] independently per voxel. Rician noise was added at signal-to-noise ratio of 35, representative of our preprocessed b = 2500 acquisition. Synthetic data were arranged in a 20 × 30 × 1 grid and fit with the identical Beltrami pipeline used on the in-vivo data (MD-based initialization per Eq. 5 of Golub et al., 2021 12, with MD_ref = 0.52 µm$^2$/ms; 200 iterations of regularized gradient descent at learning rate 0.0005; Beltrami spatial regularization weight = 1 active over the first 100 iterations). Recovered free water fractions were compared against ground truth by Pearson correlation, root-mean-square error, mean bias, and the slope of an ordinary least-squares fit. The Beltrami pipeline recovered free water fraction with high accuracy under this protocol ($r = 0.96$, RMSE = 0.078, bias = −0.024, slope = 0.92, n = 600 voxels; Supplementary Figure S1B), comparable to recovery benchmarks reported for other multi-compartment models on similar single-shell acquisitions. As expected for the single-shell regime, recovery degraded modestly at very low ground-truth FW values, where Rician noise can drive estimated MD below the tissue prior and pin the recovered FW at the lower boundary. Tissue FA recovery showed similar Pearson agreement ($r = 0.84$) but a flatter regression slope (0.77), reflecting the partial trade-off between the tissue tensor and FW compartment during fitting that is intrinsic to FWE-DTI on single-shell data.

**S2.2 Age composition of the EFW biotype**

Because free water rises physiologically at both ends of the lifespan, we examined whether EFW biotype membership was driven by the oldest participants. In both the global and regional solutions, the age of the EFW biotype was compared with that of the non-EFW biotype using Welch's unequal-variances t-test and the Mann-Whitney U test. Within the EFW biotype, the age distribution was summarized by the mean, standard deviation, median, and the number of members younger than 18 years. As a sensitivity analysis, age and sex were added as clustering features and the resulting partition was compared with the original assignment using the adjusted Rand index (ARI).

#### S2.3 Group Difference Statistical Analysis

Biotype differences in neuroimaging features and cognitive outcomes were estimated via ANCOVA. Covariates for all analyses were age and sex; TIV was additionally covaried for structural CSF volume comparisons (ventricle/TIV and extraventricular CSF/TIV) to account for residual head-size variance after normalization. Partial Cohen's d was computed as $d = t \times \sqrt{(n_1+n_2)/(n_1 \times n_2)}$. All cognitive and clinical measures were external to the clustering procedure. TD subjects were assigned post-hoc to the nearest ASD centroid (Euclidean distance in PCA space); ASD enrichment was tested via chi-square on the 2×2 contingency table.

#### S2.4 Selection of the number of clusters

To confirm that the two-cluster solution was not selected simply because $k = 2$ was the smallest partition considered, we evaluated k from 2 to 6 on the harmonized global feature set using the same procedure applied in the primary (global) analysis (k-means on the retained principal components, best of 200 initializations per k). Cluster number was assessed by the mean silhouette coefficient, and the same evaluation was applied to the regional feature set.

### S3. Supplementary Results

### S3.1 Neurocombat

Prior to harmonization, substantial site effects were present across the majority of neuroimaging features. Of 75 features, 72 (96.0%) showed significant site effects (one-way ANOVA $p < .05$), with a mean $\eta^2$ of 0.416 (range: 0.001-0.970), indicating that on average 35.8% of total variance was attributable to acquisition site rather than biology. Site effects were particularly severe for tissue axial diffusivity (ADt; mean $\eta^2 = 0.960$), tissue diffusivity metrics (MDt: mean $\eta^2 = 0.840$; FAt: mean $\eta^2 = 0.892$), and CSF/structural volumes (mean $\eta^2 = 0.662$), reflecting strong scanner- and protocol-dependent scaling differences in these measures. DTI-ALPS and FW metrics showed moderate to large site effects (mean $\eta^2 = 0.341$ and 0.274 respectively).

Following neuroCombat harmonization, site effects were eliminated across all 75 features. Zero of 75 features retained a significant site effect post-harmonization (mean $\eta^2 = 0.002$, range: <0.001-0.014), representing a 99.5% reduction in mean site effect variance. The full per-feature harmonization statistics are reported in Supplementary Tables S1 and S2.

### S3.2 ASD vs TD Group Differences

Across the full sample (N = 278), ASD participants showed significantly elevated free water (global: $d = +0.305$, $p = .012$*; FW medial: $d = +0.290$, $p = .017$*; FW posterior: $d = +0.247$, $p = .043$*), elevated tissue mean diffusivity (MDt global: $d = +0.281$, $p = .021$*), and reduced tissue fractional anisotropy (FAt WM: $d = -0.251$, $p = .039$*) relative to TD, after covarying age and sex (Supplementary Table S8). DTI-ALPS global, ALPS A/M/P, ventricular/TIV, and ReHo GM did not differ significantly (all $p > .12$). After FDR correction within each metric family, FW global ($q = .035$) and MDt global ($q = .052$) survived as marginal trends. These modest whole-sample effects are consistent with the EFW biotype (~10% of ASD) driving the group signal, while the majority of ASD participants show near-typical diffusion profiles.

#### S3.3 MRI-Cognition Regressions within ASD

Simple linear regressions (age and sex covaried, predictors standardized) within ASD ($n = 125$-$142$ depending on outcome) revealed two specific MRI-cognition associations (Supplementary Table S9). DTI-ALPS was inversely associated with FIQ ($\beta = -3.73$ per SD, $p = .009$**; ALPS medial: $\beta = -4.05$, $p = .004$**), explaining approximately 9% of IQ variance. In a multiple regression including FW, ALPS, and MDt simultaneously, ALPS remained the unique predictor of FIQ ($\beta = -3.90$, $p = .007$**). Elevated FW was associated with higher SRS Motivation scores ($\beta = +2.87$, $p = .014$*; FW medial: $\beta = +2.92$, $p = .013$*), with FW remaining the unique predictor of SRS Motivation in multiple regression ($\beta = +3.95$, $p = .021$*; FW + ALPS + MDt model). Within-biotype analyses indicate this FW-SRS Motivation association is concentrated in the EFW biotype tail: the within-EFW correlation was $r = +0.51$ ($p = .076$, $n = 13$ of 14 with SRS available), compared to $r = +0.11$ ($p = .237$, $n = 112$) in the non-EFW biotype, consistent with EFW members anchoring the high end of both distributions rather than a uniform linear relationship across all ASD. FAt WM was negatively associated with SRS Awareness ($\beta = -3.36$, $p = .003$).

#### S3.4 Age composition of the EFW biotype

In the global solution, the EFW biotype ($n = 14$) did not differ in age from the non-EFW biotype (EFW mean 21.8 years, SD 19.0, median 11.0; non-EFW mean 16.0 years, SD 12.8, median 13.0; Welch $t = 1.09$, $p = .30$; Mann-Whitney $p = .63$). The median age of the EFW biotype was in fact lower than that of the non-EFW biotype, and most EFW members were children or adolescents (8 of the 13 with available age data were younger than 18); the higher mean reflects a small number of older members rather than a systematic skew. The regional solution gave the same result (EFW $n = 15$, mean 20.7 years, median 10.5; Welch $t = 0.91$, $p = .38$). Including age and sex as down-weighted clustering features left assignments essentially unchanged (global ARI = 0.954; regional ARI = 1.000). The biotype is therefore not attributable to age: it includes participants across the age range and is not concentrated in older individuals.

#### S3.5 Selection and Number of Clusters

The mean silhouette coefficient was maximal at k = 2 and fell sharply at k = 3 and k = 4 before rising modestly at higher k (Figure S2). This profile indicates genuine two-group structure rather than a continuum being dichotomized or a finer partition being missed: had the imaging features varied continuously without latent group structure, the silhouette coefficient would not peak at k = 2 and then decline. The regional feature set showed the same k = 2 optimum (Figure S2).

**Supplementary References**

1. Di Martino A, O'Connor D, Chen B, Alaerts K, Anderson JS, Assaf M, et al. Enhancing studies of the connectome in autism using the autism brain imaging data exchange II. Sci Data. 2017;4:170010.

2. Jenkinson M, Beckmann CF, Behrens TE, Woolrich MW, Smith SM. Fsl. Neuroimage. 2012;62:782–790.

3. Fortin JP, Parker D, Tunç B, Watanabe T, Elliott MA, Ruparel K, et al. Harmonization of multi-site diffusion tensor imaging data. Neuroimage. 2017;161:149–170.

4. Fortin J-P, Cullen N, Sheline YI, Taylor WD, Aselcioglu I, Cook PA, et al. Harmonization of cortical thickness measurements across scanners and sites. Neuroimage. 2018;167:104–120.

5. Zhang Y, Brady M, Smith S. Segmentation of brain MR images through a hidden Markov random field model and the expectation-maximization algorithm. IEEE Trans Med Imaging. 2001;20:45–57.

6. Taoka T, Fukusumi A, Miyasaka T, Kawai H, Nakane T, Kichikawa K, et al. Structure of the medullary veins of the cerebral hemisphere and related disorders. Radiographics. 2017;37:281–297.

7. Virtanen P, Gommers R, Oliphant TE, Haberland M, Reddy T, Cournapeau D, et al. SciPy 1.0: fundamental algorithms for scientific computing in Python. Nat Methods. 2020;17:261–272.

8. Golub M, Neto Henriques R, Gouveia Nunes R. Free-water DTI estimates from single b-value data might seem plausible but must be interpreted with care. Magn Reson Med. 2021;85:2537–2551.

9. Jensen JH, Helpern JA. MRI quantification of non-Gaussian water diffusion by kurtosis analysis. NMR Biomed. 2010;23:698–710.

### Supplementary Tables

**Table S1. neuroCombat Harmonization: Site Effects Before and After — Summary by Metric Family**

| Metric family | Features (n) | Mean $\eta^2$ pre-ComBat | Mean $\eta^2$ post-ComBat |
|---|---|---|---|
| DTI-ALPS | 22 | 0.341 | 0.001 |
| Free water (FW) | 4 | 0.274 | 0.012 |
| Mean diffusivity (MDt) | 4 | 0.801 | 0.001 |
| Fractional anisotropy (FAt) | 4 | 0.892 | 0.006 |
| Radial diffusivity (RDt) | 4 | 0.782 | 0.003 |
| Axial diffusivity (ADt) | 4 | 0.961 | 0.002 |
| CSF / ventricular volumes | 3 | 0.662 | 0.001 |
| ReHo | 10 | 0.233 | 0.001 |
| ALFF | 10 | 0.388 | 0.001 |
| fALFF | 10 | 0.066 | 0.000 |
| **All features** | **75** | **0.416** | **0.002** |

One-way ANOVA site effects ($\eta^2$ = SS_site / SS_total) across 105 neuroimaging features grouped by metric family. N = 278 (6 sites). Sig. site = number of features with significant site effect ($p < .05$) before and after ComBat. Post-ComBat: 0/75 features retained significant site effect. Note. $\eta^2$ values: mean across features within family; range in parentheses. ADt = tissue axial diffusivity. MDt = tissue mean diffusivity (free-water corrected). FAt = tissue fractional anisotropy (free-water corrected). ReHo = regional homogeneity. ComBat model: empirical Bayes, parametric, location+scale correction; biological covariates (group, age, sex) protected.

**Table S2. Feature Selection: Drop-One Analysis — Global Feature Set (n = 7)**

| Feature | Sil (full set) | Sil (feature removed) | Δ Sil | Decision | Rationale |
|---|---|---|---|---|---|
| ***Retained features*** | | | | | |
| *DTI-ALPS global* | 0.396 | 0.421 | +0.025 | Retained | Theoretical primacy as periventricular diffusion metric; retained despite marginal Δ sil |
| *FW global mean* | 0.396 | 0.306 | **−0.090** | Retained† | Largest contributor; primary feature defining the EFW biotype |
| *MDt global mean* | 0.396 | 0.339 | **−0.057** | Retained† | Collinear with FW (r=+0.651); captures shared tissue diffusivity signal |
| *Ventricle vol/TIV* | 0.396 | 0.317 | **−0.079** | Retained† | Independent structural compartment; TIV-normalized (raw r = 0.617 with TIV) |
| *Extravent. CSF/TIV* | 0.396 | 0.332 | **−0.064** | Retained† | Extraventricular CSF independently quantified; TIV-normalized |
| *FAt WM total* | 0.396 | 0.427 | +0.031 | Retained | White matter integrity complement to MDt; theoretical completeness |
| *ReHo GM mean* | 0.396 | 0.427 | +0.031 | Retained | Exploratory functional metric; independent of diffusion features (all \|r\| < 0.13); retained but not part of the primary interpretation |
| ***Excluded features*** | | | | | |
| *ALFF GM mean* | 0.396 | 0.342 | −0.054 | Excluded | Redundant with ReHo |
| *fALFF GM mean* | 0.396 | 0.355 | −0.041 | Excluded | Reduces silhouette; adds noise without signal |

K-means (k = 2, 200 seeds, best silhouette) on z-scored PCA-reduced features (80% variance; ASD n = 149). Sil (full set) = 0.396. Δ Sil = change in silhouette when feature is removed (retained features) or added (excluded features). † = core feature (Δ sil < −0.05 when removed). DTI-ALPS = diffusion tensor image analysis along the perivascular space, treated here as a composite, structure-dependent diffusion metric rather than a direct measure of fluid clearance. FW = free water fraction. MDt = tissue mean diffusivity (free-water-corrected). FAt = tissue fractional anisotropy (free-water-corrected). TIV = total intracranial volume. ALFF = amplitude of low-frequency fluctuations. fALFF = fractional ALFF. All features ComBat-harmonized prior to analysis.

**Table S3. Feature Selection: Drop-One Analysis — Regional Feature Set (n = 17)**

| *Feature* | *Sil (full set)* | *Sil (feature removed)* | *Δ Sil* | *Decision* | *Rationale* |
|---|---|---|---|---|---|
| ***Retained features*** | | | | | |
| *ALPS anterior* | 0.391 | 0.437 | +0.046 | Retained | Theoretical; anterior medullary vein territory; ALPS position retained en bloc |
| *ALPS medial* | 0.391 | 0.428 | +0.036 | Retained | Most informative ALPS position (d=−0.79** in EFW biotype); retained en bloc |
| *ALPS posterior* | 0.391 | 0.431 | +0.040 | Retained | Theoretical; posterior medullary territory; retained en bloc with A/M set |
| *FW anterior mean* | 0.391 | 0.633 | +0.242 | Retained* | Silhouette improves without; retained for positional completeness and theoretical parity with M/P |
| *FW medial mean* | 0.391 | 0.306 | −0.085 | Retained† | Second-largest contributor; strong EFW signal (d=+3.48**) |
| *FW posterior mean* | 0.391 | 0.318 | −0.073 | Retained† | Core contributor; largest EFW signal (d=+3.56**) |
| *MDt anterior mean* | 0.391 | 0.388 | −0.003 | Retained | Marginal; retained for positional completeness across all diffusion metrics |
| *MDt medial mean* | 0.391 | 0.375 | −0.016 | Retained | Moderate contributor; elevated in EFW (d=+2.47**) |
| *MDt posterior mean* | 0.391 | 0.362 | −0.029 | Retained | Consistent contributor; elevated in EFW (d=+2.21**) |
| *FAt WM anterior* | 0.391 | 0.400 | +0.009 | Retained | Marginal; retained for positional completeness |
| *FAt WM medial* | 0.391 | 0.402 | +0.011 | Retained | Marginal; retained for positional completeness |

| | | | | | |
|---|---|---|---|---|---|
| *FAt WM posterior* | 0.391 | 0.398 | +0.007 | Retained | Marginal; retained for positional completeness |
| *ReHo GM anterior* | 0.391 | 0.409 | +0.018 | Retained | Marginal; exploratory functional metric retained en bloc with M/P |
| *ReHo GM medial* | 0.391 | 0.412 | +0.021 | Retained | Marginal; exploratory, retained en bloc |
| *ReHo GM posterior* | 0.391 | 0.404 | +0.013 | Retained | Marginal; exploratory, retained en bloc; EFW trend ($d=-0.55$, $p=.053$) |
| *Ventricle vol/TIV* | 0.391 | 0.400 | +0.009 | Retained | Structural compartment; same justification as global set |
| *Extravent. CSF/TIV* | 0.391 | 0.412 | +0.021 | Retained | Independent CSF compartment; same justification as global set |

Same procedure as Table S2. Sil (full set) = 0.391. All 17 features retained; rationale provided for features with $\Delta$ sil > 0 (improvement when removed). * = FW anterior retained for positional completeness despite $\Delta$ sil = +0.242; see rationale. † = core feature ($\Delta$ sil < −0.05 when removed).

A/M/P = anterior/medial/posterior positional means. All features derived from the same periventricular ROI parcellation as the DTI-ALPS index. Positional completeness was prioritized to maintain a full metric × position factorial design. FW anterior retained despite marginal drop-one benefit because EFW biotype membership was stable (ARI = 1.000) across sensitivity analyses.

**Table S4. Pairwise Pearson Correlations: Global Feature Set (n = 7)**

| *Feature* | *DTI-ALPS* | *FW* | *MDt* | *FAt* | *Vent/TIV* | *CSF/TIV* | *ReHo* |
|---|---|---|---|---|---|---|---|
| *DTI-ALPS global* | **+1.000** | −0.234 | −0.149 | +0.194 | −0.057 | +0.138 | −0.029 |
| *FW global mean* | −0.234 | **+1.000** | **+0.651** | −0.184 | +0.199 | −0.081 | −0.072 |
| *MDt global mean* | −0.149 | **+0.651** | **+1.000** | −0.576 | +0.105 | −0.088 | −0.009 |
| *FAt WM total* | +0.194 | −0.184 | −0.576 | **+1.000** | −0.008 | +0.084 | +0.134 |
| *Ventricle/TIV* | −0.057 | +0.199 | +0.105 | −0.008 | **+1.000** | +0.162 | −0.030 |
| *Extravent. CSF/TIV* | +0.138 | −0.081 | −0.088 | +0.084 | +0.162 | **+1.000** | +0.041 |
| *ReHo GM mean* | −0.029 | −0.072 | −0.009 | +0.134 | −0.030 | +0.041 | **+1.000** |

Computed in ASD participants (n = 149) on ComBat-harmonized, TIV-normalized features. Bold italic values: |r| ≥ 0.50. Diagonal (shaded): self-correlation. Highest collinearities: FW-MDt (r = +0.651), MDt-FAt (r = −0.576). All other pairs |r| ≤ 0.20. PCA was applied prior to clustering to mitigate residual collinearity.

**Supplementary Table S5. Pairwise Pearson Correlations: Regional Feature Set (n = 17)**

| *Feature* | *ALPS-A* | *ALPS-M* | *ALPS-P* | *FW-A* | *FW-M* | *FW-P* | *MDt-A* | *MDt-M* | *MDt-P* | *FAt-A* | *FAt-M* | *FAt-P* | *ReHo-A* | *ReHo-M* | *ReHo-P* | *Vent/TIV* | *CSF/TIV* |
|---|---|---|---|---|---|---|---|---|---|---|---|---|---|---|---|---|---|
| *ALPS anterior* | — | +0.49 | +0.19 | -0.12 | -0.15 | -0.10 | -0.17 | -0.12 | -0.04 | +0.10 | +0.06 | +0.14 | +0.02 | +0.04 | +0.14 | -0.15 | -0.00 |
| *ALPS medial* | +0.49 | — | +0.49 | -0.26 | -0.29 | -0.14 | -0.34 | -0.29 | -0.10 | +0.27 | +0.30 | +0.37 | -0.06 | -0.18 | +0.04 | -0.01 | +0.16 |
| *ALPS posterior* | +0.19 | +0.49 | — | -0.14 | -0.16 | -0.08 | -0.08 | -0.06 | +0.04 | +0.04 | +0.04 | +0.12 | -0.08 | -0.17 | +0.07 | -0.02 | +0.17 |

| | | | | | | | | | | | | | | | | | |
|---|---|---|---|---|---|---|---|---|---|---|---|---|---|---|---|---|---|
| *FW anterior* | -0.12 | -0.26 | -0.14 | — | **+0.58** | +0.40 | +0.49 | +0.38 | +0.15 | -0.11 | -0.15 | -0.16 | -0.03 | +0.04 | -0.06 | +0.11 | -0.03 |
| *FW medial* | -0.15 | -0.29 | -0.16 | **+0.58** | — | **+0.69** | +0.48 | **+0.69** | +0.40 | -0.13 | -0.13 | -0.20 | -0.06 | +0.07 | -0.04 | +0.10 | -0.13 |
| *FW posterior* | -0.10 | -0.14 | -0.08 | +0.40 | **+0.69** | — | +0.18 | +0.45 | **+0.73** | -0.09 | -0.07 | -0.18 | +0.02 | +0.02 | -0.11 | +0.24 | -0.06 |
| *MDt anterior* | -0.17 | -0.34 | -0.08 | +0.49 | +0.48 | +0.18 | — | **+0.77** | +0.32 | **-0.59** | **-0.61** | **-0.61** | -0.19 | +0.04 | -0.07 | +0.03 | -0.08 |
| *MDt medial* | -0.12 | -0.29 | -0.06 | +0.38 | **+0.69** | +0.45 | **+0.77** | — | **+0.61** | **-0.50** | **-0.54** | **-0.56** | -0.13 | +0.11 | -0.01 | +0.02 | -0.11 |
| *MDt posterior* | -0.04 | -0.10 | +0.04 | +0.15 | +0.40 | **+0.73** | +0.32 | **+0.61** | — | -0.31 | -0.34 | -0.44 | +0.06 | +0.08 | -0.04 | +0.15 | -0.06 |
| *FAt anterior* | +0.10 | +0.27 | +0.04 | -0.11 | -0.13 | -0.09 | **-0.59** | **-0.50** | -0.31 | — | **+0.89** | **+0.79** | +0.28 | -0.09 | +0.06 | -0.00 | +0.08 |
| *FAt medial* | +0.06 | +0.30 | +0.04 | -0.15 | -0.13 | -0.07 | **-0.61** | **-0.54** | -0.34 | **+0.89** | — | **+0.91** | +0.18 | -0.09 | +0.12 | +0.03 | +0.09 |
| *FAt posterior* | +0.14 | +0.37 | +0.12 | -0.16 | -0.20 | -0.18 | **-0.61** | **-0.56** | -0.44 | **+0.79** | **+0.91** | — | +0.11 | -0.07 | +0.17 | -0.05 | +0.07 |

| | | | | | | | | | | | | | | | | | |
|---|---|---|---|---|---|---|---|---|---|---|---|---|---|---|---|---|---|
| *ReHo anterior* | +0.02 | -0.06 | -0.08 | -0.03 | -0.06 | +0.02 | -0.19 | -0.13 | +0.06 | +0.28 | +0.18 | +0.11 | — | -0.01 | +0.19 | +0.09 | +0.06 |
| *ReHo medial* | +0.04 | -0.18 | -0.17 | +0.04 | +0.07 | +0.02 | +0.04 | +0.11 | +0.08 | -0.09 | -0.09 | -0.07 | -0.01 | — | -0.10 | -0.09 | +0.01 |
| *ReHo posterior* | +0.14 | +0.04 | +0.07 | -0.06 | -0.04 | -0.11 | -0.07 | -0.01 | -0.04 | +0.06 | +0.12 | +0.17 | +0.19 | -0.10 | — | -0.09 | -0.01 |
| *Ventricle/TIV* | -0.15 | -0.01 | -0.02 | +0.11 | +0.10 | +0.24 | +0.03 | +0.02 | +0.15 | -0.00 | +0.03 | -0.05 | +0.09 | -0.09 | -0.09 | — | +0.16 |
| *Extravent. CSF/TIV* | -0.00 | +0.16 | +0.17 | -0.03 | -0.13 | -0.06 | -0.08 | -0.11 | -0.06 | +0.08 | +0.09 | +0.07 | +0.06 | +0.01 | -0.01 | +0.16 | — |

Computed in ASD participants (n = 149). Bold italic values: |r| ≥ 0.50. Diagonal (—): self-correlation. Abbreviations: A = anterior, M = medial, P = posterior. High within-metric collinearities: FAt-A-FAt-M (r = +0.887), FAt-M-FAt-P (r = +0.907), MDt-A-MDt-M (r = +0.769), and MDt-FAt cross-metric pairs (r = −0.50 to −0.61). These within-metric positional correlations are expected; PCA dimensionality reduction (8 PCs, 83.2% variance) resolves collinearity prior to clustering, retaining independent variance components.

**Table S6.** Cluster Validity Metrics Across Algorithms and Feature Sets

| ***Feature set*** | ***Algorithm*** | ***k*** | ***Silhouette*** | ***Perm p*** | ***Null mean ± SD*** | ***ARI vs k-means***[a] |
|---|---|---|---|---|---|---|
| ***Global (5 PCs, 87.4%)*** | k-means ★ | 2 | 0.396 | <.0001 | −0.002 ± 0.060 | — |
| | GMM (BIC) | 2 | 0.383 | — | — | 0.954 |
| | Hierarchical | 2 | 0.381 | — | — | 0.906 |
| ***Regional (8 PCs, 83.2%)*** | k-means ★ | 2 | 0.391 | <.0001 | −0.003 ± 0.058 | — |
| | GMM (BIC) | 3 | 0.339 | — | — | 0.615 |
| | Hierarchical | 2 | 0.391 | — | — | 1.000 |

★ = primary solution. Perm p = permutation p-value (n = 2,000). GMM k selected by Bayesian Information Criterion (BIC). Hierarchical = Ward linkage. PCs = principal components at ≥ 80% variance.

**Cluster-number selection across k**

**Table S7.** TD Validation: Nearest-Centroid Projection of TD Subjects into ASD-Derived Clusters

| | ***EFW biotype*** | ***Non-EFW cluster*** | ***Enrichment test*** | | |
|---|---|---|---|---|---|
| | ASD | TD (projected) | ASD | TD (projected) | |
| ***Global analysis (7 features, 5 PCs)*** | | | | | |
| *n (%)* | 14 (9.4%) | 6 (4.7%) | 135 (90.6%) | 123 (95.3%) | $\chi^2$ = 1.675, p = .196 (ns) |
| *ASD proportion* | 70.0% | — | 52.3% | — | — |
| *FW global mean* | 0.447 ± 0.039 | 0.381 ± 0.027 | 0.343 ± 0.018 | 0.347 ± 0.019 | — |
| *MDt global mean* | 0.00081 ± 0.00010 | 0.00072 ± 0.00003 | 0.00063 ± 0.00003 | 0.00063 ± 0.00003 | — |
| *DTI-ALPS global* | 1.075 ± 0.104 | 1.099 ± 0.079 | 1.148 ± 0.093 | 1.152 ± 0.095 | — |
| ***Regional analysis (17 features, 8 PCs)*** | | | | | |
| n (%) | 15 (10.1%) | 7 (5.4%) | 134 (89.9%) | 122 (94.6%) | $\chi^2$ = 1.456, p = .228 (ns) |
| *ASD Proportion* | 68.2% | — | 52.3% | — | — |
| FW posterior mean | 0.485 ± 0.068 | 0.397 ± 0.041 | 0.358 ± 0.028 | 0.362 ± 0.025 | — |
| MDt medial mean | 0.00074 ± 0.00011 | 0.00068 ± 0.00004 | 0.00062 ± 0.00004 | 0.00062 ± 0.00003 | — |
| ALPS medial | 1.086 ± 0.142 | 1.163 ± 0.094 | 1.193 ± 0.121 | 1.196 ± 0.113 | — |

TD subjects assigned post-hoc to nearest ASD centroid (Euclidean distance in PCA space). Values are mean ± SD. EFW = elevated free-water biotype. TD subjects played no role in clustering. Enrichment test: chi-square on 2×2 contingency (biotype × group). ASD proportion = ASD/(ASD+TD) within cluster. **p < .01; ns = not significant.

**Table S8. ASD vs TD Group Differences in Neuroimaging Markers**

| ***Feature*** | ***ASD mean ± SD*** | ***TD mean ± SD*** | ***d (partial)*** | ***p*** | ***q (FDR)*** |
|---|---|---|---|---|---|
| ***Diffusion MRI — FW (ANCOVA: age + sex)*** | | | | | |
| *FW global mean* | 0.354 ± 0.038 | 0.346 ± 0.024 | +0.305* | .012 | 0.035*† |

| | | | | | |
|---|---|---|---|---|---|
| *FW anterior mean* | 0.330 ± 0.040 | 0.324 ± 0.026 | +0.232† | .056 | 0.056† |
| *FW medial mean* | 0.347 ± 0.040 | 0.338 ± 0.027 | +0.290* | .017 | 0.035*† |
| *FW posterior mean* | 0.361 ± 0.044 | 0.351 ± 0.029 | +0.247* | .043 | 0.056† |
| ***Diffusion MRI — MDt, FAt, ALPS (ANCOVA: age + sex)*** | | | | | |
| *MDt global mean (mm²/s)* | 6.4e-4 ± 0.4e-4 | 6.3e-4 ± 0.3e-4 | +0.281* | .021 | 0.052†† |
| *FAt WM total* | 0.362 ± 0.033 | 0.368 ± 0.028 | −0.251* | .039 | 0.056† |
| *DTI-ALPS global* | 1.145 ± 0.096 | 1.150 ± 0.079 | −0.085 | .485 | 0.578 |
| *ALPS medial* | 1.186 ± 0.123 | 1.202 ± 0.098 | −0.190 | .119 | 0.477 |
| *ALPS anterior* | 1.148 ± 0.108 | 1.148 ± 0.094 | −0.007 | .956 | 0.956 |
| *ALPS posterior* | 1.099 ± 0.122 | 1.100 ± 0.107 | −0.015 | .903 | 0.956 |
| ***Structural volumes (ANCOVA: age + sex + TIV)*** | | | | | |
| *Ventricle vol / TIV* | 0.0569 ± 0.0097 | 0.0557 ± 0.0082 | +0.087 | .477 | 0.578 |
| *Extravent. CSF / TIV* | 0.0447 ± 0.0059 | 0.0444 ± 0.0057 | +0.003 | .978 | 0.956 |
| ***Resting-state fMRI (ANCOVA: age + sex)*** | | | | | |
| *ReHo GM mean* | 0.128 ± 0.087 | 0.135 ± 0.075 | −0.095 | .433 | 0.630 |

Partial Cohen's d from ANCOVA (age + sex covaried; additionally TIV for structural volumes). FDR: Benjamini-Hochberg applied within each metric family separately (FW x4, MDt x4, FAt x4, ALPS x4, CSF x2, ReHo x4). *N = 278 (ASD n = 149, TD n = 129)*. FW = free water fraction. MDt = tissue mean diffusivity. FAt = tissue fractional anisotropy. TIV = total intracranial volume. CSF volumes TIV-normalized and winsorized at ±3 SD prior to analysis. FDR applied within each metric family: FW (4 tests), MDt (4 tests), FAt (4 tests), ALPS (4 tests), CSF volumes (2 tests), ReHo (4 tests). *p < .05; +p < .10 (uncorrected). *q < .05; +q < .10 (FDR within metric family).

### Table S9. MRI-Cognition Regressions within ASD

| *MRI predictor (z-scored)* | *β* | *p* | *q (FDR)* | *R²* | *n* |
|---|---|---|---|---|---|
| ***Outcome: Full-Scale IQ (covariates: age, sex)*** | | | | | |
| *DTI-ALPS global* | −3.73 | .009$^{**}$ | 0.100$^{\dagger}$ | 0.086 | 141 |
| *ALPS medial* | −4.05 | .004$^{**}$ | 0.073$^{\dagger}$ | 0.094 | 141 |
| *MDt global mean* | −0.26 | .862 | 0.922 | 0.032 | 142 |
| *FAt WM total* | −0.20 | .896 | 0.922 | 0.032 | 142 |
| *FW global mean* | −0.37 | .802 | 0.922 | 0.032 | 142 |
| *Ventricle vol / TIV* | −0.87 | .567 | 0.827 | 0.034 | 139 |
| ***Outcome: SRS Total T-score (covariates: age, sex) — n = 125*** | | | | | |
| *FW global mean* | +2.39 | .031$^{*}$ | 0.157 | 0.090 | 125 |
| *FW medial mean* | +2.49 | .026$^{*}$ | 0.150 | 0.093 | 125 |
| *MDt global mean* | +2.07 | .062$^{\dagger}$ | 0.212 | 0.082 | 125 |
| *FAt WM total* | −1.93 | .086$^{\dagger}$ | 0.250 | 0.077 | 125 |
| *DTI-ALPS global* | +0.27 | .809 | 0.922 | 0.055 | 125 |
| *ALPS medial* | −0.34 | .762 | 0.922 | 0.055 | 125 |
| ***Outcome: SRS Motivation T-score (covariates: age, sex) — n = 125*** | | | | | |
| *FW global mean* | +2.87 | .014$^{*}$ | 0.100$^{\dagger}$ | 0.052 | 125 |
| *FW medial mean* | +2.92 | .013$^{*}$ | 0.100$^{\dagger}$ | 0.053 | 125 |
| *MDt global mean* | +1.54 | .189 | 0.441 | 0.017 | 125 |
| *DTI-ALPS global* | +0.74 | .533 | 0.827 | 0.006 | 125 |
| *ALPS medial* | +0.71 | .549 | 0.827 | 0.006 | 125 |
| ***Outcome: SRS Awareness T-score (covariates: age, sex) — n = 125*** | | | | | |
| *FAt WM total* | −3.36 | .003$^{**}$ | 0.073$^{\dagger}$ | 0.085 | 125 |
| *ALPS medial* | −2.26 | .047$^{*}$ | 0.196 | 0.049 | 125 |
| *MDt global mean* | +2.23 | .050$^{\dagger}$ | 0.196 | 0.048 | 125 |
| *FW global mean* | +1.11 | .334 | 0.616 | 0.025 | 125 |
| ***Outcome: ADOS-G Total (covariates: age, sex) — n = 88*** | | | | | |
| *FW global mean* | +0.02 | .964 | 0.964 | 0.012 | 88 |

| | | | | | |
|---|---|---|---|---|---|
| *DTI-ALPS global* | +0.28 | .521 | 0.827 | 0.019 | 87 |
| *MDt global mean* | +0.10 | .820 | 0.922 | 0.013 | 88 |
| *FAt WM total* | −0.46 | .296 | 0.616 | 0.025 | 88 |

Simple linear regressions, one standardized MRI predictor at a time (covariates: age + sex). q = FDR-corrected p-value across all predictor-outcome pairs (BH correction, 35 tests total). Only imaging features with $p < .10$ for a given outcome are shown; ADOS-G shown for completeness. β = unstandardized regression coefficient per 1 SD of predictor. $R^2$ = full model R-squared (including covariates). All analyses within ASD (n = 149; n varies by outcome due to missing data).

**Table S10. PC1 Loadings: Regional Feature Set (n = 17)**

| Feature | PC1 loading | Domain |
|---|---|---|
| MDt medial | +0.384 | White matter microstructure |
| MDt anterior | +0.366 | White matter microstructure |
| FAt WM posterior | -0.361 | White matter microstructure |
| FAt WM medial | -0.343 | White matter microstructure |
| FAt WM anterior | -0.327 | White matter microstructure |
| FW medial | +0.288 | Extracellular fluid |
| MDt posterior | +0.279 | White matter microstructure |
| FW posterior | +0.237 | Extracellular fluid |
| DTI-ALPS medial | -0.227 | Periventricular directional diffusivity |
| FW anterior | +0.223 | Extracellular fluid |
| DTI-ALPS anterior | -0.118 | Periventricular directional diffusivity |
| DTI-ALPS posterior | -0.091 | Periventricular directional diffusivity |
| ReHo GM anterior | -0.079 | Local functional coherence |
| ReHo GM posterior | -0.070 | Local functional coherence |

| Feature | PC1 loading | Domain |
|---|---|---|
| ReHo GM medial | +0.065 | Local functional coherence |
| Ventricle vol / TIV | +0.064 | CSF compartment |
| Extraventricular CSF / TIV | -0.059 | CSF compartment |

Loadings on the first principal component of the 17-feature regional set, fit on z-scored features imputed to feature-wise ASD medians across all 149 ASD participants. PC1 explained 28.9% of total variance. Sign was oriented so that positive loadings indicate a greater multivariate imaging deviation. Features are ordered by absolute loading. PC1 correlated with FW global mean at r = +0.67 ($R^2 = 0.44$) within the ASD group, indicating that it captures MDt and FAt variance partially independent of free water alone. FW = free water; MDt = tissue mean diffusivity; FAt = tissue fractional anisotropy; ReHo = regional homogeneity; TIV = total intracranial volume.

**Table S11. PC1 Partial Correlations with Clinical Measures within ASD**

| Outcome | n | partial r | p | q (FDR) |
|---|---|---|---|---|
| SRS Awareness | 125 | +0.253 | 0.005 | 0.026 |
| SRS Cognition | 125 | +0.201 | 0.026 | 0.046 |
| SRS Communication | 125 | +0.199 | 0.028 | 0.046 |
| SRS Motivation | 125 | +0.125 | 0.170 | 0.212 |
| SRS Mannerisms | 125 | +0.083 | 0.364 | 0.364 |
| SRS Total | 125 | +0.195 | 0.031 | — |
| ADOS-G Total | 88 | +0.062 | 0.573 | — |
| Full-Scale IQ | 142 | +0.029 | 0.730 | — |

Partial correlations between PC1 and each clinical outcome within the ASD group, residualized for age and sex by ordinary least squares. Benjamini-Hochberg FDR was applied across the five SRS treatment subscales (k = 5); SRS Total, ADOS-G Total and Full-Scale IQ are shown for completeness and were not included in the correction family. PC1 = first principal component of the 17-feature regional set; SRS = Social Responsiveness Scale; ADOS-G = Autism Diagnostic Observation Schedule-Generic.

**p < .01; *p < .05; †p < .10 (uncorrected). †q = FDR q < .10.

### Supplementary Figures

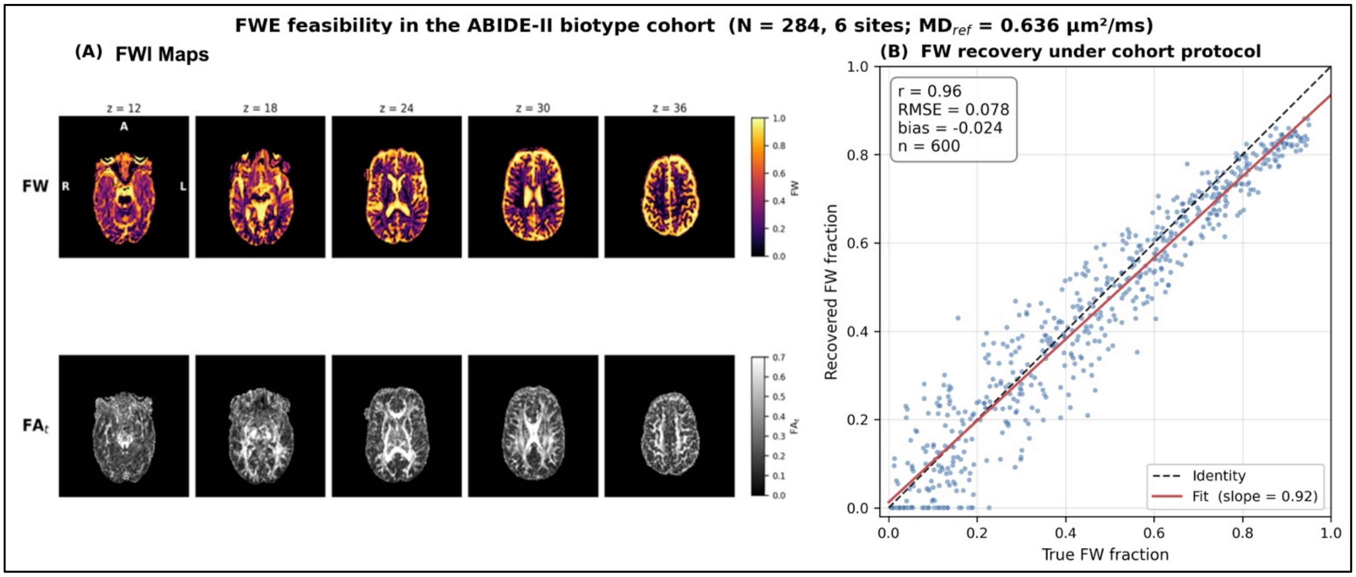


**Figure S1. FWE feasibility in the ABIDE single-shell data.** (A) Beltrami FWE compartment maps from one participant computed with the Golub et al. (2021) implementation: free water fraction (FW, top) and FW-corrected tissue fractional anisotropy ($FA_t$, bottom) shown across five axial slices spanning ventricular and supraventricular levels. Orientation: R/L = right/left, A = anterior. (B) FW recovery under the ABIDE protocol: signals synthesized from known free water fractions and fit with the same Beltrami pipeline applied to *in vivo* data track ground truth with strong rank preservation (r = 0.96, RMSE = 0.078, bias = −0.024, slope = 0.92, n = 600 voxels).

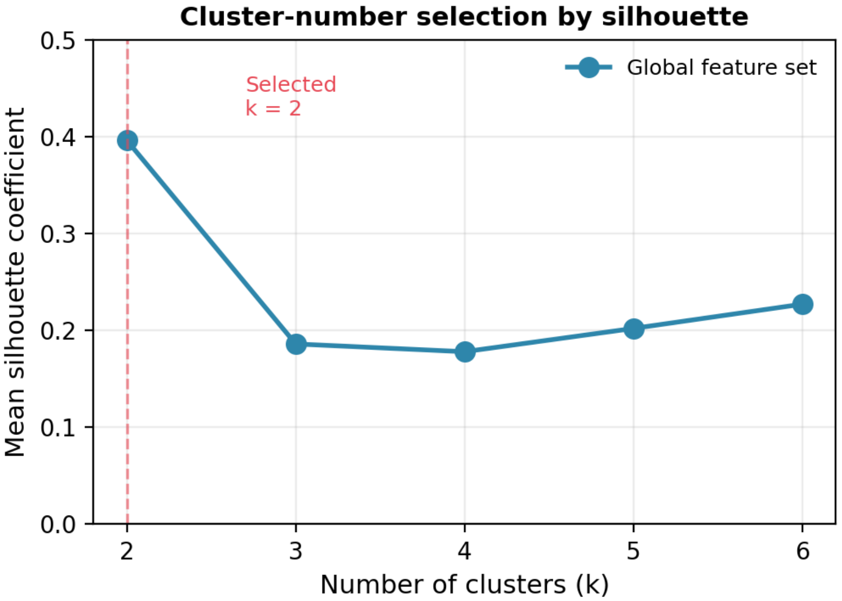


**Figure S2. Mean silhouette coefficient as a function of the number of clusters k for the harmonized global feature set.** The coefficient peaks at k = 2 and declines at k = 3 to 4, supporting the two-cluster solution. k = 2 value as reported in Table S6 (0.396); values for k = 3 to 6 computed with the identical pipeline (neuroCombat-harmonized global features, k-means on 5 principal components, best of 200 initializations). The sharp decline from k = 2 to k = 3 supports the two-cluster solution. k = 2 silhouette = 0.396; k = 3 silhouette = 0.186; k = 4 silhouette = 0.178; k = 5 silhouette = 0.202; k = 6 silhouette = 0.227.
